# Defective synapto-nuclear signaling contributes to motoneuron vulnerability in SOD1-ALS

**DOI:** 10.64898/2026.09.01.748568

**Authors:** Kamil Grycz, Cédric Jan, Piotr Zawistowski, Bartosz Wasicki, Burak Özkan, Oumayma Aousji, Riccardo Sirtori, Claudia Fallini, Guillaume Caron, Simon M. Danner, Francesco Roselli, Daniel Zytnicki, Marcin Bączyk

## Abstract

Glutamatergic excitatory synapses not only shape spiking activity and neuronal communication but also initiate activity-dependent signaling pathways that trigger transcriptional programs. Since glutamatergic excitatory synapses onto spinal motoneurons (MNs) are impaired presymptomatically in Amyotrophic Lateral Sclerosis, we investigated whether synapto-nuclear coupling is disrupted in MNs from mSOD1 mice and whether restoring it mitigates pathology. We developed an *in vivo* approach to selectively investigate the coupling between synaptic excitation and nuclear CREB phosphorylation in spinal MNs. Specific activation of Ia-MN synapses induces CREB phosphorylation in wild-type MNs but not in mSOD1 MNs at P50, indicating presymptomatic synapto-nuclear uncoupling. Enhancing cAMP/PKA signaling by pharmacological inhibition of cAMP degradation restored synapto-nuclear coupling, reduced misfolded SOD1, and slowed neuromuscular-junction denervation. Thus, activity-dependent synapto-nuclear signaling is impaired yet pharmacologically rescuable in mSOD1 MNs, supporting synapto-nuclear signaling as a determinant of MN resilience.

## Introduction

Synapses are traditionally conceptualized for their ability to shape spiking activity and neuronal communications. Yet, synaptic activity also drives signaling from active synapses to the nucleus to shape activity-dependent transcriptional programs involved in synaptic homeostasis [1], metabolic adaptation [2], plasticity [3], and neuronal resilience to insults [4].

Multiple signaling pathways are set in motion at postsynaptic sites by Ca^2+^ influx through NMDAR, AMPAR, and Voltage-Dependent Calcium Channels [5]. In particular, Ca^2+^-sensitive adenylcyclases raise cAMP levels upon synaptic activity, [6] freeing the catalytic subunit of PKA from the regulatory/inhibitory subunit and allowing PKA phosphorylation of cytoplasmic and nuclear substrates [7]. In parallel, Ca^2+^/Calmodulin complexes activate CamKII and CaMKIV; CamKII primarily mediates local signaling, whereas CaMKIV can migrate to the nucleus and regulate gene transcription [8, 9]. Furthermore, Erk cascade is also activated at the synaptic level by glutamate receptors through RAS-GRF proteins, providing another route from synaptic activity to nuclear responses [10, 11].

Synapse-activated signaling pathways converge on the transcription factor CREB [12], which contains multiple consensus phosphorylation sites for PKA, Erk, Rsk [13] and CaMKIV [5]. Phosphorylated CREB (pCREB) drives transcriptional programs that support neur,onal survival (CREB loss leads to widespread neuronal apoptosis [5], synaptic plasticity, mitochondrial biogenesis, and neuromuscular junctions (NMJ) maintenance [14–17]). Synaptic signaling also regulates protein synthesis through the Erk/RSK/mTOR pathway [18]; phosphorylation of ribosomal proteins is an outcome of synaptic activity and indirectly represents neuronal firing[19].

Due to its impact on gene modules critical for neuronal resilience and survival, altered synaptic signaling is hypothesized to influence neuronal vulnerability to disease [20, 21]. In Amyotrophic Lateral Sclerosis (ALS), multiple lines of evidence suggest that synaptic inputs are disturbed in the early stages of the disease. In the mSOD1 mouse model, synaptically evoked excitatory postsynaptic potentials (EPSP) [22] are reduced in amplitude, matching the loss of PSD-95, Homer, Shank proteins, and synaptic glutamate receptors [22–24]. The appearance of synaptic alterations is shared also by multiple iPSC-generated models with mutations in C9ORF72, TPD-43 and CHCHD10 mutations, either at transcriptome/proteome level [25–27]. or at neuron/network level [28]. Human tissue studies likewise reveal synaptic alterations by histopathology [29] or proteomics [25]. Consistent with these findings, disturbances in the nuclear targets of synapto-nuclear signaling have been repeatedly demonstrated, in particular on CREB and SRF [30, 31].

Given that glutamatergic excitatory synapses are impaired in ALS, we asked whether synapto-nuclear coupling is disrupted in motoneurons (MNs) of mSOD1 mice and sought to identify the steps at which this pathway fails. We then tested whether synapto-nuclear signaling can be restored and if its restoration reduces disease marker burden. Such findings would implicate defective synapto-nuclear signaling as a contributor to ALS pathophysiology.

Here we exploited the arrangement of Ia proprioceptive inputs to MN, namely the unique dynamic sensitivity of spindle primary endings to vibrations and the distinct histochemical profile of Ia-MN synapses (large, convoluted vGluT1+ terminals [32, 33]), to systematically probe synapse-to-nucleus signal transduction evoked by stereotyped synaptic activation and to probe small-molecule interventions at multiple levels to restore synapto-nuclear signaling. We found that: 1) activity-dependent synapto-nuclear signaling is uncoupled in MNs from the mSOD1 mice, and 2) a boost of the cAMP/PKA pathway, achieved by pharmacological inhibition of cAMP degradation, restores the synapto-nuclear coupling and reduces disease-marker burden.

## Results

### Synapto-nuclear signaling from Ia afferents in spinal motoneurons

We developed an *in vivo* protocol in wild-type (WT) mice to investigate and isolate the coupling between Ia synapse excitation and signaling in spinal MN (Figure 1). The key step is a specific activation of spindle primary endings in the triceps surae muscle (TS) with high frequency vibration of the Achilles tendon (400 Hz, 0.6 mm peak-to-peak in anesthetized mice; Fig. 1B, Supplemental Fig. 1 and Methods). This allows selective activation of the Ia-MN synapses on TS of the “vibrated side”, since Ia afferents from hindlimb muscles do not project to the contralateral side [34]. *In vivo* intracellular recordings confirmed that such vibrations induced Ia excitatory post- synaptic potentials (EPSPs) in TS-innervating MNs (Supplemental Figure 1), and that these EPSPs remained subthreshold. Thus, any activation of downstream pathways in our preparation can primarily be ascribed to synaptic activity rather than action potential firing.

**Figure 1:**
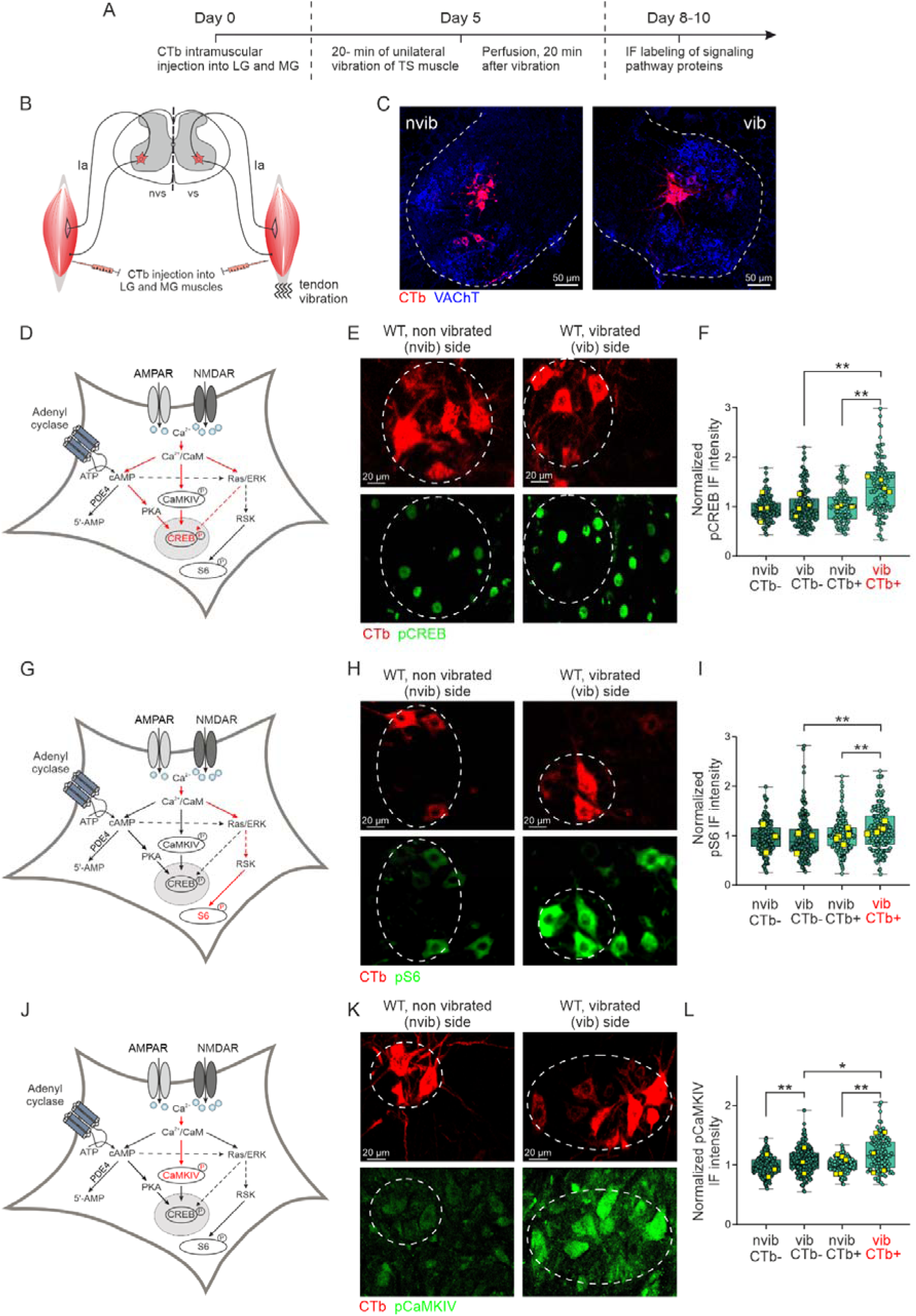
Functional activation of the Ia-MN synapses engages the activity-dependent signaling pathways in TS MNs from WT mice. **A,** experimental design for investigating the activity-dependent pathways in WT mouse spinal MNs. **B,** graphical schematic showing bilateral retrograde labelling of triceps surae (TS) MNs and unilateral Achilles tendon vibration. **C,** Labelled CTb MN, showing the Triceps surae pool, on both sides of the spinal cord (images from the same section). **D,** intracellular signaling pathways (red) responsible for the activity-dependent CREB phosphorylation. Upon activation of the Ia-MN synapses, calcium enters the cell through the NMDA and AMPA receptors, activating the calcium/calmodulin complex and then three pathways (cAMP/PKA, CamKIV, Ras/ERK) that converge to phosphorylate CREB and S6 (the later missing the CamKIV component). **E,** representative images of pCREB labelling in CTb-labelled TS MNs from the vibrated and non-vibrated side of the lumbar spinal cord. **F,** boxplots displaying the distribution of fluorescence intensity of pCREB in CTb+ and CTb- MNs of both vibrated and non-vibrated sides. Notice a significantly 46% stronger normalized pCREB signal in CTb+ MN from the vibrated side (1.46±0.19) in comparison to both CTb+ MN from the non-vibrated (1.00±0.10, estimated marginal means (EMMs) ratio=1.37±0.07, t_477_=6.424, p<0.01, LMM on log-transformed outcomes) and CTb- MNs from the vibrated side (0.99±0.15; EMMs ratio=1.41±0.06, t_469_=7.827, p<0.01, LMM on log-transformed outcomes). Furthermore, the interaction effect is significant confirming that the effect of vibration was stronger in CTb+ than CTb- MNs (EMMs ratio=1.38±0.09, F_1,476.9_=24.797, p<0.01; LMM on log-transform outcomes; n=4 mice). **G**, intracellular signaling pathways (red) responsible for the activity-dependent S6 phosphorylation. **H,I**, representative images and data quantification for pS6 labeling. Similarly to pCREB, a significant increase in normalized FI can be seen both in CTb+ MNs of the vibrated side (1.16±0.4) in comparison to both CTb+ MNs form the non-vibrated side (1.00±0.05; EMMs ratio=1.09±0.03, t_740_=2.818, p<0.01, LMM on log-transform outcomes) and CTb- MNs from the vibrated side (0.91±0.13; EMMs ratio=1.21±0.04, t_65_=5.233, p<0.01, LMM on log-transform outcomes). Furthermore, the interaction effect was significant indicating that vibration had a stronger effect in CTb+ MNs than in CTb- ones (EMMs ratio=1.2±0.05; F_1,729.6_=15.491, p<0.01; LMM on log-transform outcomes; n=6 mice). **J-L,** similar organization as for D-F and G-I but showing pCaMKIV data. Again a significant increase in normalized FI signal is present for CTb+ MNs of the vibrated side (1.19±0.14) compared to the non-vibrated side (1.00±0.07; EMMs ratio=1.16±0.03, t_671_=6.502, p<0.01, LMM on log-transform outcomes) and compared to CTb- MNs from the vibrated side (1.04±0.12; EMMs ratio=1.06±0.03, t_279_=2.315, p=0.02, LMM on log-transform outcomes). Furthermore, the interaction effect was significant indicating that indeed vibration had a stronger effect on FI signals of CTb+ than CTb- MNs (EMMs ratio=1.06±0.03, F_1,671.6_=4.601, p=0.03; LMM on log-transform outcomes; n=5 mice). In addition, significantly higher CaMKIV levels can be identified in CTb- MNs of the vibrated side in comparison to CTb- MNs of the non-vibrated side (1.04±0.21 vs. 0.95±0.19 for CTb- MNs of the vibrated and non-vibrated side, respectively, EMMs ratio =1.09±0.02, t671=4.788, p<0.01, MM on log-transform outcomes, n=5 mice). In the boxplots, each dot indicates a single MN, whereas the large squares represent the average values for each mouse. The lower and upper hinges correspond to the first and third quartiles (25th and 75th percentiles, respectively). The upper whisker extends from the hinge to the largest value while the lower whisker extends from the hinge to the smallest value. The horizontal line on the boxplots represents the median of the whole group. The significant differences between the investigated groups are marked as * for p<0.05, ** for p<0.01. Scale bars: 20 µm, the dashed ovals indicate the analysed MN pools, as guided by CTb labelling. LMM - Linear Mixed Model, GLMM - Generalized Linear Mixed Model, EMMs - estimated marginal means.

Our protocol started with bilateral injections of CTb in TS muscles in order to retrogradely label the corresponding motoneuron pools (Figure 1B,C). Five days later, the right (test) TS received 20-min tendon vibration, followed by a 20-min post-stimulation interval to allow for the signaling cascade to be fully activated. The mouse was then euthanized by intracardial perfusion. Upon section immunolabelling for phosphorylated CREB (pCREB) on Ser133, we revealed that the vibration sequence induced a significant increase of pCREB in CTb+ MNs (MNs belonging to the TS pool) on the stimulated side compared with CTb+ MNs on the non-stimulated side (+46% on average; Figure 1D-F). In sharp contrast, the vibration did not upregulate pCREB in CTb- MNs (non-TS MNs expected to receive little or no heteronymous Ia synaptic excitation; Figure 1D-F). Vibration-specific phosphorylation increases were also observed for the ribosomal protein S6 (+16% on average), suggesting that the Ca^2+^/CaM Ras/ERK pathway was active (Figure 1G-I), and for CAMKIV (+19% on average), suggesting some contribution of Ca^2+^/CaM CAMKIV (Figure 1J-L). We conclude that our vibration protocol engages multiple activity-dependent signaling pathways selectively within the MN pool that receives homonymous Ia input from the stimulated muscles.

### The synapto-nuclear signaling is dysfunctional in MN from mSOD1 mice

Next, we applied the vibration protocol/phosphoprotein staining assay to probe synapto-nuclear signaling in mSOD1 mice (Figure 2). In the mSOD1 mice, vibration-induced Ia activation did not elicit the CREB phosphorylation response, in sharp contrast to WT mice (Figure 2 B-D). Similar failures in phosphorylation increase were observed for pS6 (Figure 2E-G) and pCAMKIV (Figure 2H-J). Together, these data indicate that synaptic activity-dependent signaling is impaired in the MNs from mSOD1 mice. This impairment could reflect reduced representation of glutamatergic receptors at VGluT1 synapses [22], dysfunction in downstream signaling of the receptors, or a combination of the two.

**Figure 2:**
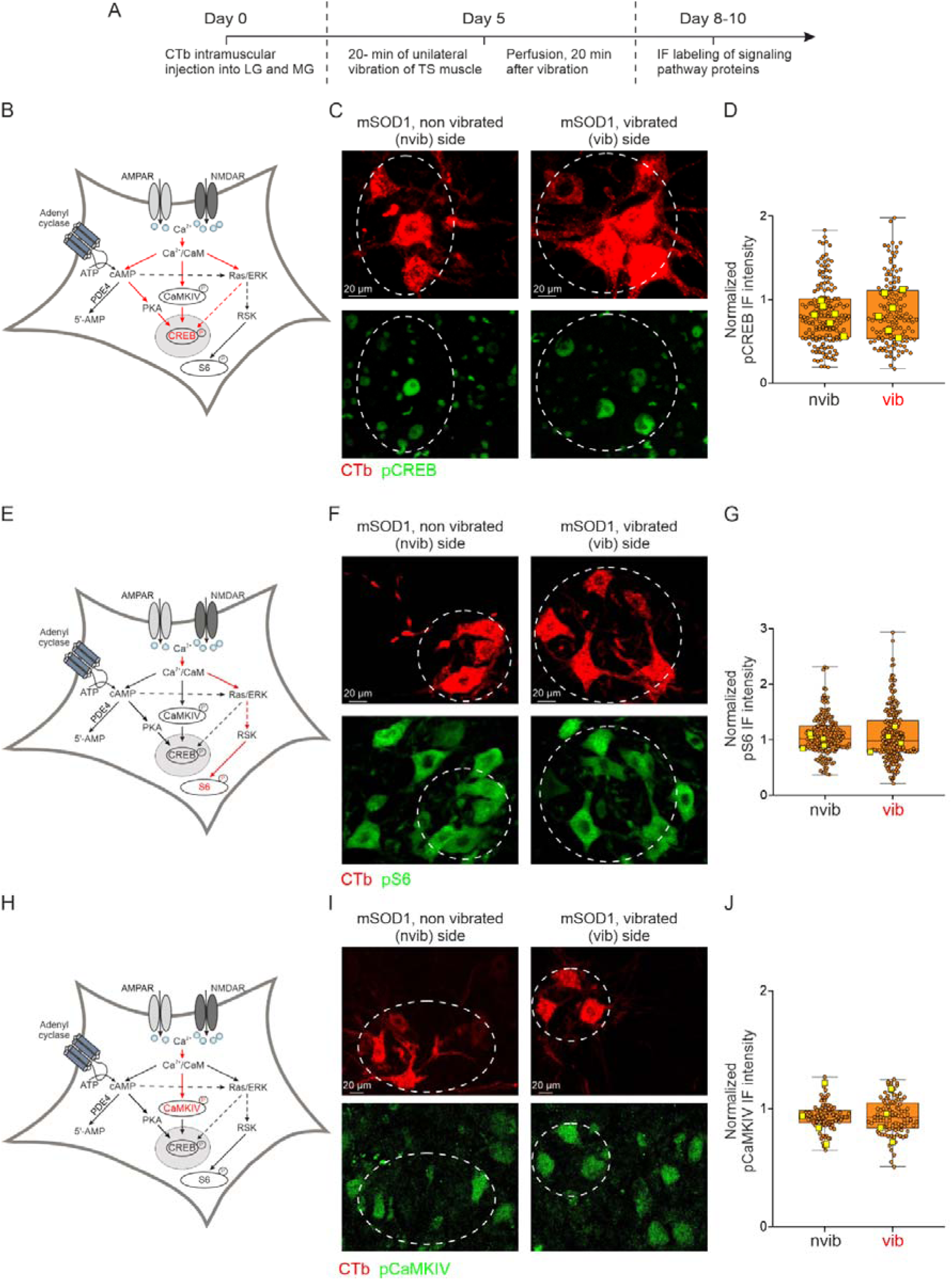
Synapse to nucleus signaling is impaired in MNs from mSOD1 mice. **A,** experimental design for investigating the activity-dependent pathways in mouse spinal MN. **B,** graphical schematic showing activity-dependent pathways converging to phosphorylate CREB enlightened in red. **C**, representative images, and **D** data distribution of normalized fluorescence intensity of pCREB in CTb-labelled neurons in the non-vibrated and vibrated side of the lumbar spinal cord. Notice a lack of increase in pCREB signal in response to vibration (0.85±0.10 for non-vibrated vs. 0.88±0.10 for vibrated side, EMMs ratio=1.03±0.12, t_764_=0.666, p=0.506, LMM on log-transforme outcomes, n=6 mice). **D - F,** and **G - I,** similar organization as for **A-C**, but for the metabolic pathway inducing S6 phosphorylation **(D)** or CamKIV phosphorylation **(G)** highlighted in red. Again, vibrations do not increase S6 phosphorylation **(E, F)** (1.00±0.05 vs. 0.97±0.0.09, EMMs ratio=0.96±0.34, t_inf_=-0.693, p=0.49, LMM on log-transformed outcomes, n=5 mice) and CamKIV phosphorylation **(H, I)** (0.93±0.11 and 0.92±0.10, EMMs ratio=1.00±0.03, t_860_=-0.044, p=0.97, LMM on log-transformed outcomes, n=4 mice) levels for MNs in the non-vibrated side compared with the vibrated side, respectively. Boxplots description as in Figure 1. Scale bars: 20 µm, the dashed ovals indicate the analysed MN pools, as guided by CTb labelling.

### Boosting AMPA receptor does not restore synapto-nuclear coupling, whereas boosting NMDA activation induces only a partial restoration

We elected to dissect the mechanisms involved in the failure of activity-dependent synaptic signaling in mSOD1 MN by first focusing on the glutamatergic receptors level. VGluT1 synapses onto MN are endowed not only with AMPA receptors [22] but also with NMDA and mGluR5 receptors (Supplemental Figure 2; in agreement with the general architecture of excitatory synapses [35]). In fact, together with reduced synaptic GluR4 AMPAR previously demonstrated [22], we now also observed a trend toward reduced representation of NMDA receptors, as the total NMDA cluster size at VGluT1 synapses tend to be smaller in MNs from mSOD1 mice than from WT mice (Supplemental Figure 2). We then asked if boosting the signaling of AMPAR or NMDAR would be sufficient to restore the coupling between vibration stimulation and pCREB/pS6 readouts.

We first attempted to restore the signaling through AMPA receptors using Ampakine CX516 which is known to prolong the opening of AMPA receptors and to increase the size and duration of EPSPs [36]. We verified that i.v. injection of Ampakine CX516 in mSOD1 mice results in the significant increase of vibration-induced EPSPs (Figure 3B-D). The EPSP amplitude increase upon Ampakine injection is likely caused by a larger synaptic current since it cannot be explained by an increase of the MN input resistance (in fact, a decrease was observed; Figure 3E), or by an hyperpolarization of the resting membrane potential (unchanged; Figure 3F). Next, we investigated if the ampakine-induced increase in EPSP amplitude translated into a restoration of downstream signaling events. To this end, Ampakine CX516 (40 mg/kg, delivered i.p., as previously reported [37, 38]) was injected 50 minutes before the vibration. To our surprise, upon vibration, Ampakine produced only a negligible increase in pCREB (6%, Figure 3 G,H) and no change in pS6 (Figure 3 I,J). This suggests that magnifying AMPAR currents does not restore the engagement of the activity-dependent downstream signaling.

**Figure 3:**
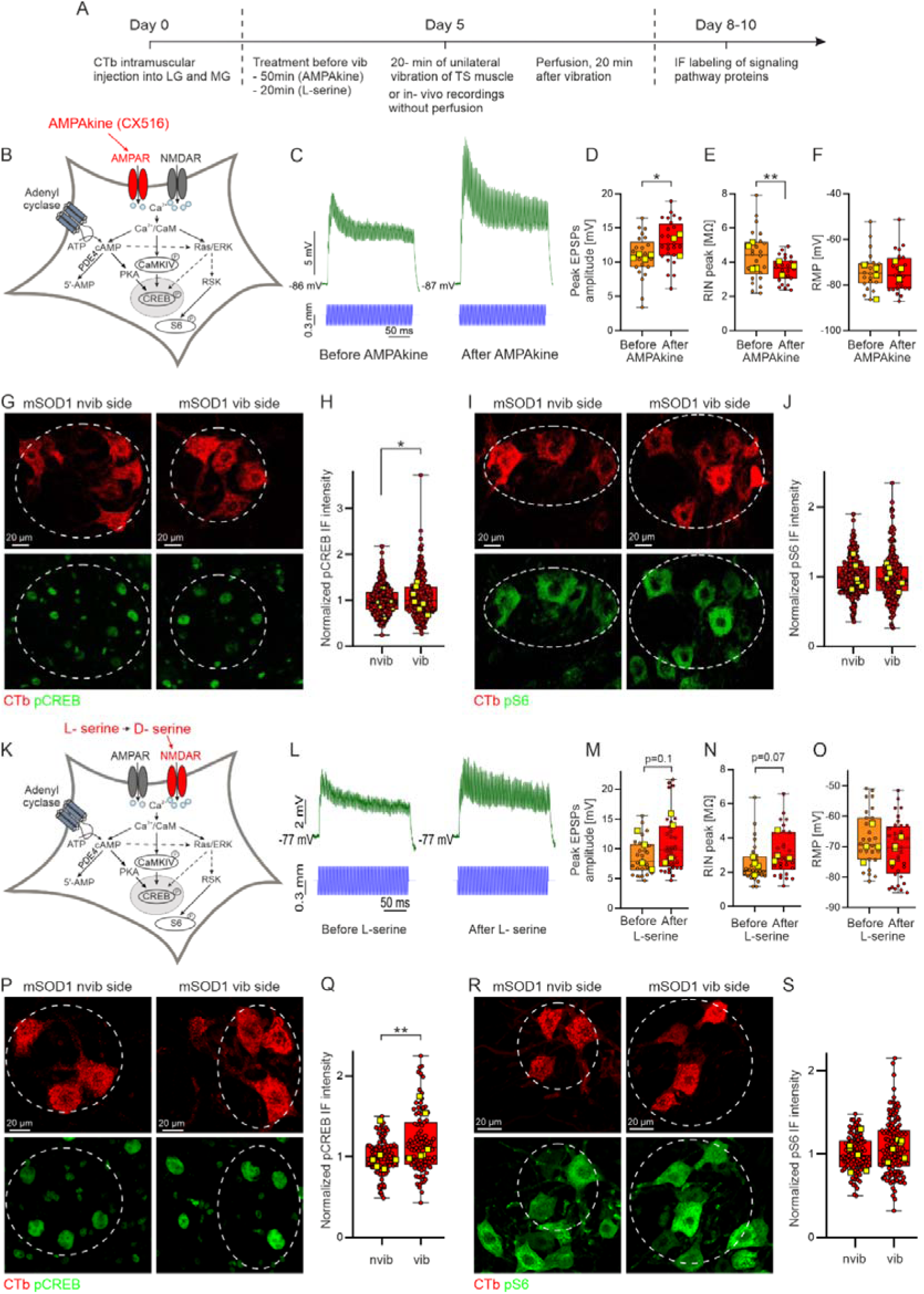
Activation of NMDA receptors, not AMPA receptors, helps to restore the activity-dependent pathways in mSOD1 G93A mice. **A,** experimental design for investigating the effects of upregulation of AMPA and NMDA receptor activity on the cAMP/PKA, CamKIV, Ras/ERK pathways in MNs of mSOD1 G93A mice. **B,** a diagram of the approach to increase AMPA receptor activity with CX516 (Ampakine). **C,** representative voltage traces of vibration-induced EPSP intracellularly recorded from two TS MNs before (left) and after (right) Ampakine treatment, top trace (green): voltage response, bottom trace (blue): vibrator arm movement. **D-F**, boxplots displaying the data distribution for compound EPSP amplitude (**D**), cell input resistance (**E**) and RMP (**F**) before vs. after the treatment. Notice a strong increase in the peak compound EPSP amplitude (from 10.81±14 mV to 12.81±0.0.69 mV, EMMs estimate 2.13±0.97, t_44_=2.203, p=0.03, LMM), followed by a decrease in input resistance (from 4.35±0.0.34 MΩ to 3.70±0.0.17 MΩ, EMMs estimate=-1.14±0.40, t_14.7_=-2.872, p=0.01, LMM, n=4 mice) and no change in the RMP (−76.55±2.78 mV vs. −73.32±1.60, EMMs estimate=-0.99±2.56, t_43_=-0.385, p=0.70, LMM, n=4 mice). **G - H,** representative images (**G**) and boxplot of data distribution (**H**) of normalized pCREB fluorescence intensity from CTb-labelled MNs from the vibrated and non-vibrated side of the lumbar spinal cord following Ampakine treatment. Notice that increased AMPA receptor activity results only in a modest 6% increase in pCREB response levels (0.97±0.09 in non-vibrated vs. 1.03±0.10 in vibrated side, EMMs estimate=- 0.01±0.23, t_612_=-0.482, p=0.63, LMM, n=7 mice). **I-J**, as for **G-H**, but showing the lack of impact of Ampakine treatment on the pS6 levels (1.01±0.08 in non-vibrated vs. 1.01±0.06 in vibrated side, EMMs estimate=0.02±0.02, t_480_=0.883, p=0.38, LMM, n=6 mice). **K-L** similar as for **B-J**, but showing impact of L-serine treatment on spinal MN electrophysiological profile (**M-O**), and the normalized pCREB and pS6 levels in MNs of vibrated vs. non-vibrated side (**P-S**). Notice the lack of significant effect of L-serine treatment on the vibration-induced EPSPs amplitudes (9.48±1.20 mV before vs. 11.5±1.65 mV after L-serine treatment, EMMs ratio=1.13±0.08, t_53.2_=1.685, p=0.10, LMM on log-transformed outcomes, n=4 mice, **M**), input resistance (2.4±0.18 MΩ before vs. 3.22±0.33 MΩ after L-serine treatment, EMMs ratio=1.24±0.15, t_50_=1.819, p=0.07, LMM on log-transformed outcomes, n=4 mice, **N**) and the resting membrane potential (−69.41±2.15 mV before vs. −70.58±1.42 mV after L-Serine treatment, EMMs estimate=-3.62±2.51, t_54.7_=-1.439, p=0.16, LMM, n=4 mice, **O**). However, L-serine treatment evokes a significant 21% increase in the vibration-induced CREB phosphorylation (1.02±0.09 in non-vibrated vs. 1.23±0.13 in vibrated side, EMMs ratio=1.15±0.45, t_8.69_=3.381, p=0.01, LMM on log-transformed outcomes, n=6 mice **P-Q**), while it similarly fails to alter the pS6 levels (1.01±0.08 in vibrated vs. 1.06±0.05 in non-vibrated side, EMMs ratio=1.06±0.45, t_9.85_=1.237, p=0.25, LMM on log-transformed outcomes, n=6 mice, **R-S**). Scale bars: 20 µm, the dashed ovals indicate the analysed MN pools, as guided by CTb labelling. Boxplot and statistical description as in Figure 1.

Then, we aimed to boost the opening of the NMDAR acting at the co-agonist level. Since D-serine barely crosses the blood-brain barrier [39], we used L-serine that crosses the blood- brain barrier more efficiently and is converted into D-serine through serine racemase enzymatic activity in about 20 minutes [40]. This treatment produced a trend (p=0.10) toward increasing the size of the vibration-induced EPSP (Figure 3 K-M), but this increase might partly be explained by an increase of the input resistance (Figure 3 N). Nevertheless, L-serine treatment amplified the vibration-elicited increase of pCREB (21%, Figure 3 P,Q) although still not affecting pS6 levels (Figure 3 R,S). Taken together these data reveal that boosting AMPAR or NMDAR opening results in a limited restoration of CREB but not of S6 phosphorylation. Thus, reduced receptor opening constitutes only a component of the disturbed synapto-nuclear signaling in mSOD1 MN.

### Boosting the cAMP/PKA pathway allows to restore the synapto-nucleus coupling

Since boosting the opening of glutamate receptors produced only an incomplete recovery of synaptic signaling, we explored the possibility of further alterations occurring downstream. Among the plethora of signaling pathways initiated by synaptic activity [5], we elected to target the cAMP/PKA pathway [41] since it is involved in the downstream signaling [5] and in the regulation of synaptic glutamate receptors [42]. We find evidence of a direct disturbance of the synaptic cAMP/PKA signaling in the mSOD1 MN. First, the synaptic clusters (in vGluT1+ synapses) of the PKA-anchoring protein (AKAP 79/150 also known as AKAP5), which concentrates PKA heterotetramer close to adenylcyclases and to its substrates [43], show a strong tendency (p=0.05) toward smaller size in mSOD1 MN (Figure 4 A-E). Second, the cytoplasmic and nuclear content of the catalytic PKA subunits are reduced in MN from mSOD1 mice (Figure 4 J-N). Furthermore, both cytoplasmic (p=0.06) and nuclear levels of the catalytic PKA subunit (Supplemental Figure 3 A-F), as well as phosphorylated CREB (Supplemental Figure 3 D,G) are reduced in cortical neurons from SOD1 patients compared with controls.

**Figure 4:**
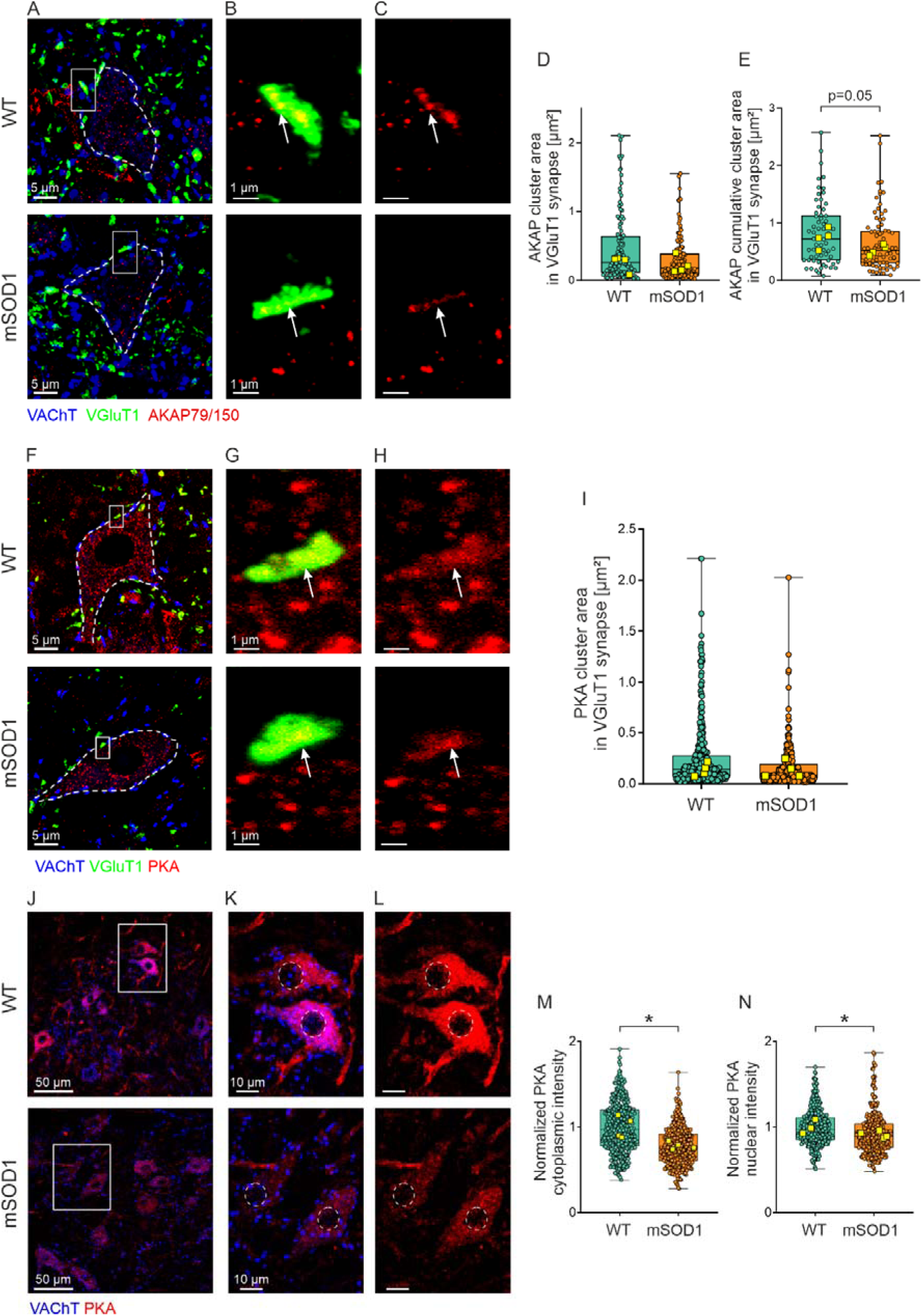
Disturbance of the synaptic cAMP/PKA signaling in MN from the mSOD1 mice. **A,** representative image of PKA-anchoring protein (AKAP79/150) clusters in TS MNs of WT (top) and mSOD1 (bottom) mice. **B-C**, magnification of the boxed region in **A**, showing a representative VGluT1 synapse with AKAP clusters marked by white arrowheads. **D-E**, boxplots showing the data distribution for AKAP cluster area (0.46±0.08 µm^2^ vs. 0.33±0.06 µm^2^, EMMs estimate=-0.18±0.35, t_5.98_=-0.521, p=0.62, LMM, n=8 mice) and AKAP cumulative cluster area (0.83±0.09 µm^2^ vs. 0.67±0.07 µm^2^, EMMs estimate=-0.26±0.13, t_139_=- 1.954, p=0.05, LMM on log-transformed scale, n=8 mice) in WT and mSOD1 mice respectively. **F-I** similar as for **A-E**, but showing the representative image **F-H**, and data distribution (**I**) for the lack of difference in catalytic PKA subunits area in the VGluT1 synapse (white arrowheads in **G-H** point to the PKA clusters) between WT (0.26±0.05 µm^2^) and mSOD1 mice (0.16±0.03 µm^2^, EMMs ratio=0.66±1.67, t_6_=-1.614, p=0.16, LMM on log-transformed scale, n=8 mice). In **A** and **B**, the MN soma is marked by dashed white lines as traced by VAChT labelling. **J-L**, representative images of WT and mSOD1 MNs labelled for cytoplasmic and PKA levels. **M, N**, boxplots displaying the normalized data distribution of lower cytoplasmic PKA (1.00±0.06 vs. 0.78±0.02, EMMs ratio=0.79±0.56, t_5.98_=-3.368, p=0.02, LMM on log-transformed outcomes, n=8 mice) and nuclear PKA (1.00±0.03 vs. 0.91±0.02, EMMs ratio=0.91±0.32, t_5.86_=-2.610, p=0.04, LMM on log-transformed scale, n=8 mice) content in WT and mSOD1 animals respectively. **K** and **L**, show the magnified regions in **J** with MNs nuclei marked with dashed white ovals. Boxplot and statistical description as in Figure 1. Scale bar 5 µm for main panels and 1 µm for magnifications.

In order to enhance the PKA signaling, we used Rolipram, an inhibitor of the phosphodiesterase-4 [44], to boost cAMP levels by preventing its degradation. The administration of a single dose of Rolipram (1 mg/kg), 40 minutes before the vibrations sequence produced no effect on the vibration-induced EPSP (Figure 5 B,C) nor on the input resistance (Figure 5 D), or the resting membrane potential (Figure 5E), indicating no interference with glutamate receptors in synapses. However, it significantly increased the pCREB level upon vibrations (+22% on average, Figure 5 F-H) and substantially increased also pS6 levels (+20% on average Figure 5 I-K). Rolipram does not increase CamKIV phosphorylation (Figure 5 L-N). Altogether, this experiment demonstrates that boosting the cAMP/PKA pathway with Rolipram is sufficient to restore an activity-dependent phosphorylation of both CREB and S6.

**Figure 5:**
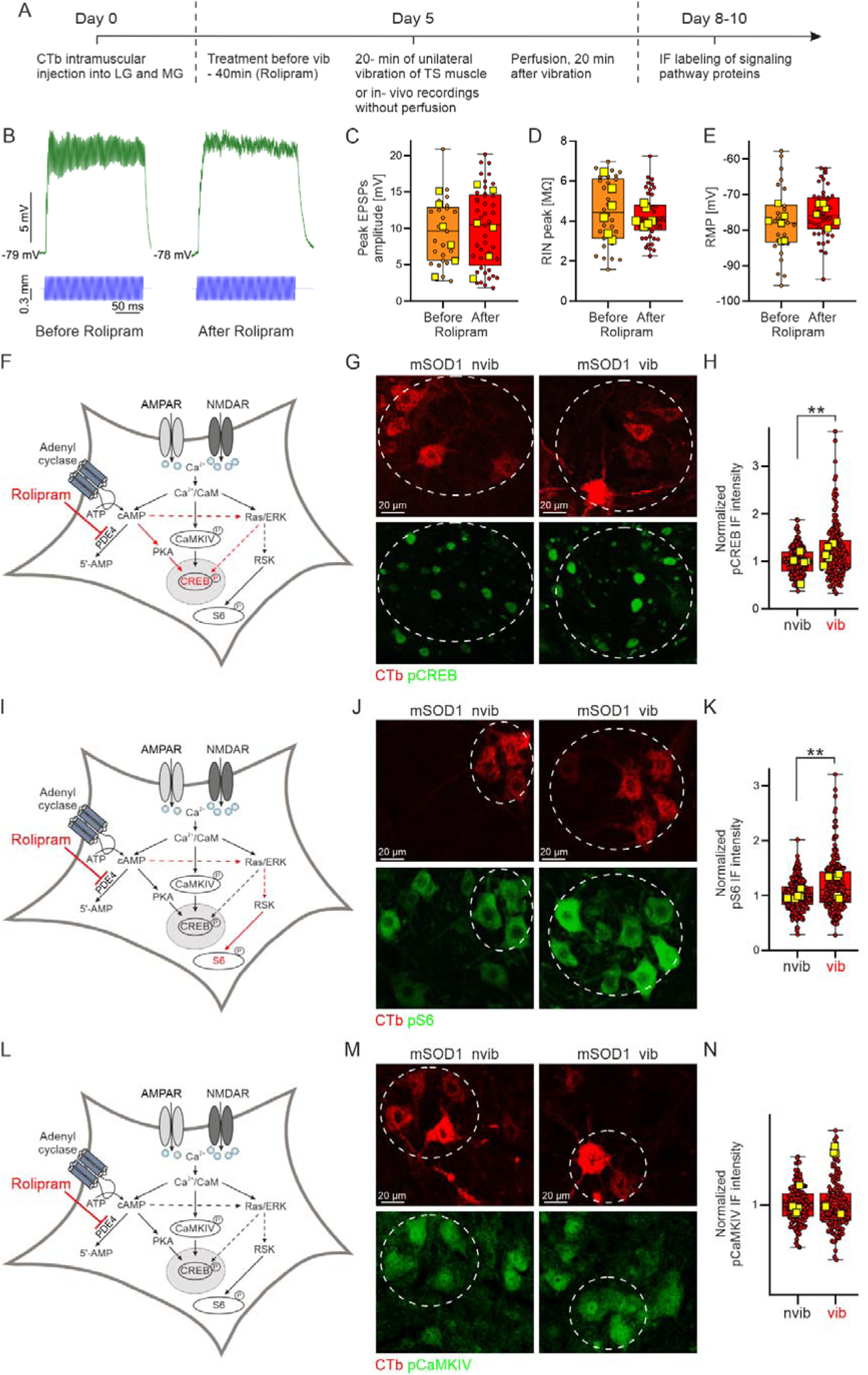
A single dose of Rolipram (a phosphodiesterase-4 inhibitor) treatment restores activity-dependent signaling in mSOD1 mice. **A**, experimental design for investigating the effects of a single dose of Rolipram on cAMP/PKA, Ras/ERK, CamKIV pathways activity in MNs of mSOD1 G93A mice. **B**, representative voltage traces of TS MN response to achilles tendon vibration before (left) and after (right) rolipram treatment. Top trace - intracellularly recorded voltage response, bottom trace - vibrator arm movement. **D-E**, Boxplots showing the data distribution for maximal EPSP amplitude (9.21±1.85 mV vs. 10.2±2.05 mV, EMMs estimate 0.91±0.90, t(_58.6_)=1.008, p=0.32, LMM, n=6 mice, D), peak input resistance (4.57±0.53 MΩ vs. 4.02±0.19 MΩ, EMMs estimate −0.32±0.34, t(_66.6_)=-0.946, p=0.35, LMM, n=6 mice, E), and resting membrane potential (−78.1±1.69 mV vs. −75.3±1.17 mV, EMMs estimate 2.12±2.18, t(_63_)=0.971, p=0.34, LMM, n=6 mice, F), for TS MNs recorded before and after Rolipram treatment respectively. Notice a lack of effect of Rolipram treatment on any of the aforementioned parameters. **F**, a diagram showing the target engagement of Rolipram treatment for cAMP/PKA pathway activation. **G-H,** representative images and boxplot of data distribution for pCREB fluorescence intensity in TS MNs of mSOD1 animals, in the vibrated and non-vibrated side. Notice a significant increase in normalized pCREB FI in MNs of the vibrated side after the Rolipram treatment (0.95±0.11 vs. 1.16±0.08, for non-vibrated vs. vibrated side respectively, EMMs estimate −0.14±0.04, t(_410_)=-3.715, p<0.01, LMM on log-transformed outcomes, n=5 mice). **I**, a diagram showing the target engagement of Rolipram treatment for Ras/ERK pathway activation. **J,** similar as for **H**, but showing representative images and data distribution for pS6 FI. Again a significant increase in pS6 FI can be seen in MNs of the vibrated side (1.00±0.03 vs. 1.20±0.10, for non-vibrated vs. vibrated side respectively, EMMs estimate −0.13±0.03, t(_437_)=-4.209, p<0.01, n=5 mice). **K** and **L** similar as for **J-H**, but showing target engagement and FI levels for Rolipram activation of the CamKIV pathway. This time, no significant effect of the treatment can be seen (1.00±0.07 vs. 1.00±0.06 for non-vibrated and vibrated side respectively, EMMs estimate 0.01±0.02, t(_281_)=0.585, p=0.56, LMM on log-transform outcomes, n=5 mice). Images scale bar 20 µm, boxplot and statistical description as in Figure 1.

### Boosting the cAMP/PKA pathway with Rolipram reduces ALS disease burden

Since a single dose of Rolipram is able to restore (at least partially) the activity-dependent CREB and S6 phosphorylation, we explored if repeated rolipram administration for 10 days (10 mg/kg per day, i.p.) had beneficial effects on ALS disease burden. Note that during these ten days, mice were freely moving in the animal facility, and many excitatory synapses onto MNs were active in order to ensure posture and locomotion; among those are Ia synapses, since muscle spindles are stretched during motor activities contributing to the proprioceptive feedback.

Consistent with the acute findings, ten-day Rolipram treatment did not significantly affect the size of vibration-induced EPSPs, input resistance, or resting membrane potential (Figure 6 C-E), excluding the contribution of synaptic modification. However, repeated injections of Rolipram increased baseline (i.e., without vibration) pCREB levels (+43% on average, p=0.05 Figure 6 F,G) and pS6 (+27% on average) (Figure 6 H,I), confirming successful engagement of cAMP/PKA-dependent targets. Since restoration of pCREB is associated with neuroprotection [30], we further explored the impact of Rolipram on disease markers. Rolipram reduced the burden of misfolded SOD1 protein by 21% in MNs of mSOD1 mice compared with vehicle-treated animals (Figure 7 B,C). Furthermore, Rolipram treatment increased the fraction of neuromuscular junctions that remained fully innervated at the age of P50 both in Tibialis Anterior (+18% on average) and in Triceps Surae (+16% on average) (Figure 7 D-G).

**Figure 6:**
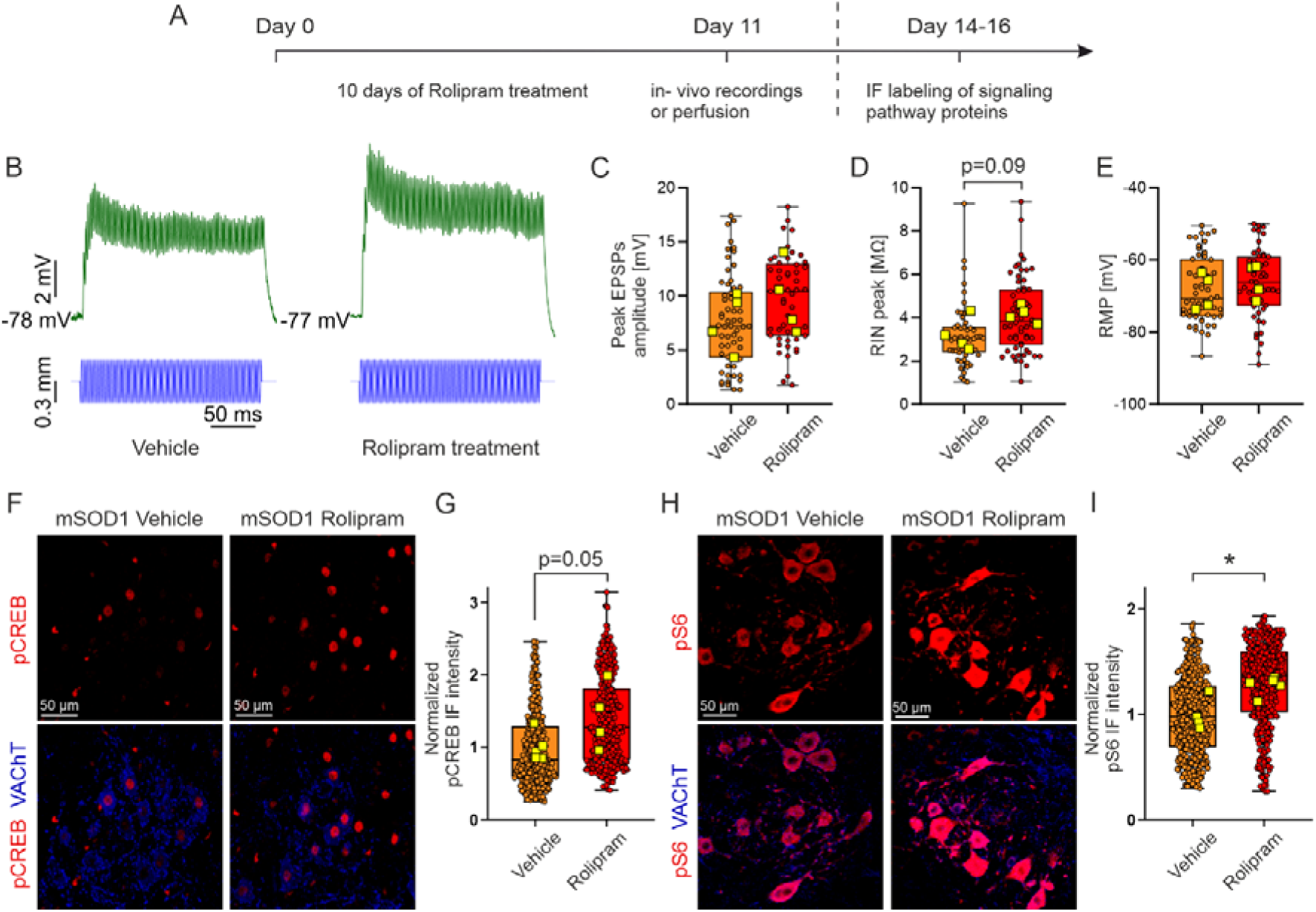
Chronic delivery of Rolipram restores the activity-dependent signaling in mSOD1 mice. **A**, experimental design for investigating the effects of chronic rolipram treatment on MNs electrophysiological profile and signaling proteins in mSOD1 mice. **B**, Intracellular recordings of voltage traces from TS MN in response to Achilles tendon vibration in the Vehicle (left) and Rolipram-treated (right) group. Top trace - intracellularly recorded voltage response, bottom trace - vibrator arm movement. **C**, boxplots presenting the data distribution for TS MNs electrophysiological profile in the Vehicle and Rolipram-treated groups. No significant effect of the treatment can be identified on maximal EPSP (**C**, 7.69±1.34 mV vs. 9.8±1.64 mV, EMMs estimate=2.25±2.21, t_5.87_=1.020, p=0.35, LMM, n=4 mice), peak input resistance (**D,** 3.23±0.39 MΩ vs. 4.17±0.20 MΩ, EMMs ratio=1.29±0.16, t_5.2_=2.025, p=0.09, LMM on log-transformed outcomes, n=4 mice), and resting membrane potential (**E,** −68.77±2.52 mV vs. −65.86±2.34 mV, EMMs estimate=2.08±3.53, t_5.5_=0.589, p=0.58, LMM, n=4 mice) for the Vehicle vs. Rolipram-treated group, respectively. **F-G**, representative images and boxplots displaying a significantly higher normalized pCREB fluorescence intensity in TS MNs from Rolipram-treated in comparison to Vehicle-treated animals (1,00±0.09 vs. 1.43±0.22, for non-treated vs. treated animals, respectively, EMMs ratio=0.68±0.12, t_6.98_=-2.286, p=0.05, LMM on log-transformed outomes, data n=10 mice). **H-I**, similar as in F-G but showing representative images and data distribution of increase of normalized pS6 FI from 1.00±0.08 in the Vehicle to 1.27±0.04 in the Rolipram-treated group (EMMs estimate=-519±155, t_6.95_=-3.358, p=0.01, LMM, n=10 animals). Boxplot and panel description as in Figure 1.

**Figure 7:**
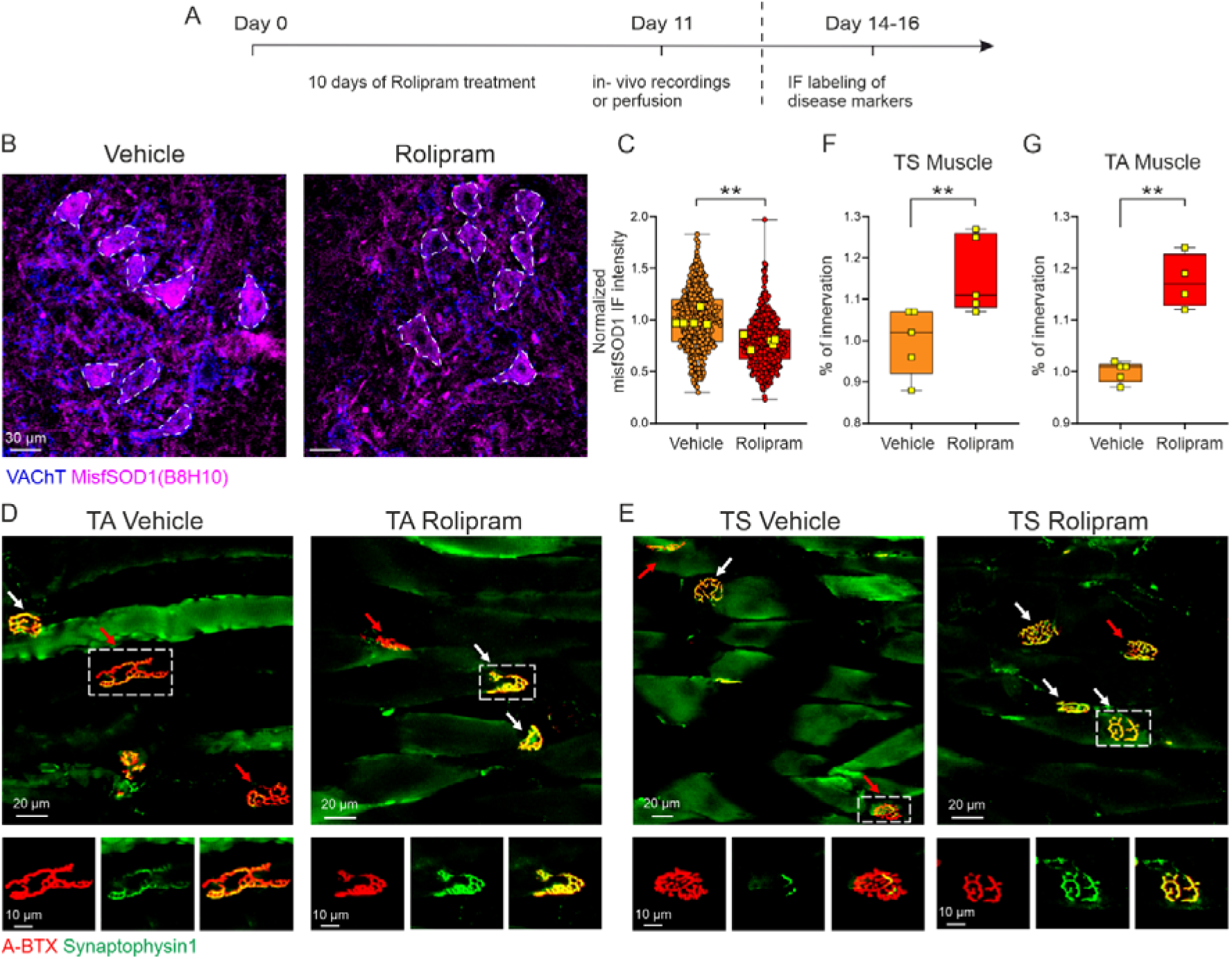
Chronic delivery of Rolipram reduces misfolded SOD1 protein and protects neuromuscular junction from denervation. A, experimental design for investigating the effects of chronic rolipram treatment on disease markers and neuromuscular junction condition in mSOD1 mice. **B-C**, representative images and boxplots, showing a reduction of misfolded SOD1 protein levels in the rolipram treated group (normalized data 1.00±0.03 vs. 0.79±0.03, in vehicle and rolipram treated groups respectively, estimate=-197±40.3, t_7.97_=-4.891, p<0.01, LMM, n=10 mice). **D-G**, representative images and boxplots showing the percentage of fully innervated neuromuscular junctions in the fast-contracting TA, and mixed TS muscles. A significantly higher fraction of fully innervated neuromuscular junctions can be seen following Rolipram treatment in both TA (normalized data 1.00±0.01 vs. 1.18±0.02, for Vehicle vs. Rolipram treated groups, estimate=-7.83±1.15, t_14_=6.649, p<0.01, LMM, n=10 mice) and TS (1.00±0.04 vs. 1.16±0.04, for Vehicle vs. Rolipram treated groups, estimate=-9.64±3.39, t_16_=-2.847, p=0.01, LMM, n=10 mice) muscles.

## Discussion

We developed an *in vivo* approach to investigate the coupling between synaptic excitation and nuclear CREB phosphorylation through activity-dependent signaling pathways in spinal motoneurons. Selective activation of the Ia-MN synapses produces CREB phosphorylation in MNs from WT mice but not in MNs from mSOD1 mice at P50, indicating presymptomatic synapto-nuclear uncoupling. We observed deficits consistent with impaired cAMP/PKA signaling, and showed that pharmacologic enhancement of this pathway with Rolipram (an inhibitor of the phosphodiesterase-4) partially restored synapto-nuclear coupling. Rolipram also reduced misfolded SOD1 in MNs and increased the fraction of fully innervated neuromuscular junctions.

### An approach to selectively investigate synapto-nuclear coupling in spinal MN

The excitatory synapses formed between Ia afferents, originating from the spindle primary endings in skeletal muscles, and spinal MNs are the most appropriate to investigate the synapto-nuclear coupling in MNs, due to the particular anatomo-functional organization of Ia afferents. First, spindle primary endings, innervated by Ia afferents, display a high dynamic sensitivity allowing them to respond efficiently to small muscle vibrations at high frequencies whereas secondary endings, innervated by group II afferents, have a much lower dynamic sensitivity [45]. Golgi tendon organs, innervated by Ib afferents, display little sensitivity to muscle stretches [46]. Second, among muscle, skin or joint proprioceptive afferents, only the Ia afferents make monosynaptic connections with spinal MN [47]. Third, the excitatory action of Ia afferents from TS is mostly restricted to ipsilateral ankle extensor MNs [48]. Indeed, hindlimb Ia afferents do not cross the spinal cord and thereby do not make monosynaptic connection with contralateral MNs [34]. Moreover, in anaesthetized mammals, Ia afferents from TS do not elicit synaptic actions on interneurons strong enough to make them firing and to indirectly activate MN (disynaptic EPSP are barely seen in ipsilateral (for cats [47, 48], rats [49], for mice [22]) and in contralateral triceps surae MNs [50]. Therefore, in our experimental conditions, Ia afferents, activated by the Achilles tendon vibrations, elicit a rather specific excitation of ipsilateral TS MNs. CREB, S6 and CamKIV phosphorylations induced by unilateral vibrations of TS are in keeping with the anatomo-functional organization of Ia afferents from TS (phosphorylations are mostly observed in TS MNs (CTB+) from the vibrated side, not in those from the contralateral side and not in MNs others than triceps surae (CTB-)). The CREB, S6 and CamKIV phosphorylations are then the consequence of a specific activation of the Ia-MN excitatory synapses in TS MN.

### Multiple causes for synapto-nuclear uncoupling

We demonstrated that Ia synaptic activation does not translate in CREB and S6 phosphorylation in the mSOD1 MNs, in contrast to what happens in WT animals, indicating a degree of uncoupling between Ia synaptic activity and synaptic signaling, and particularly from the nuclear signaling. Given the central role of CREB in regulating neuronal survival and overall cell biology, this uncoupling may deprive MNs of a critical neuroprotective drive and disrupt multiple domains of cell biology including axonal transport, mitochondrial metabolism [51], autophagy [52], and overall stress resilience [21].

Synapto-nuclear uncoupling may, theoretically, take place at one (or more) of the several signaling steps in the cascade. First, the synapto-nuclear signaling depends on the calcium that enters the MN in response to Ia synaptic activity (in agreement with the model established for other neurons [53]). Several sources of calcium may be involved: Ca^2+^- permeable glutamate receptors, namely NMDAR and AMPAR subtypes (in particular GluR2 with edited Q/R site [54]), and Cav1.2 and Cav1.3 voltage-dependent Ca^2+^ channels activated by the synaptically-induced depolarization [55–57]. Alterations in the synaptic representation of several AMPAR subunits were previously reported [22]. Here, we have also reported a trend in the decrease in synaptic NR1 subunits (shared by all NMDAR receptors [58]) in vGluT1+ synapses. Regarding the voltage-dependent calcium channels, even if the expression level of Cav1.3 channels were not found to be altered in MNs from mSOD1 mice, [59] we cannot exclude that the smaller Ia EPSPs in MNs from mSOD1 mice [22] might activate these channels to a lesser extent. Whatever the case, reduced signaling may originate because of the reduced number of glutamate receptors (AMPAR, NMDAR) as Ca^2+^ sources.

This model is made less likely by our findings that ampakines and possibly L-serine can potentiate the synaptic currents upon Ia activation, with only a modest restoration of the signaling cascades. Thus, boosting synaptic glutamate receptors currents may be necessary but remains not sufficient to re-establish synapto-nuclear signaling. On the other hand, limiting the degradation of cAMP by Rolipram was sufficient to reinstate PKA signaling upon vibration in keeping with the critical role of the cAMP/PKA pathway in activity-dependent CREB phosphorylation.

Interestingly, similar defects in the activation of the CREB pathway were described in TDP-43 mutant neurons [60] as well as in iPSC-derived neurons and postmortem tissue of human patients with mutations in the C9ORF72 gene [61]. suggesting that alterations to this pathway may be a common feature of multiple forms of ALS.

Since an enhancement of NMDAR and AMPAR activation does not allow a significant signaling restoration, whereas a boost of PKA produces a significant restoration of the synapto-nuclear signaling, we suggest that a critical failure point (not necessarily the only alteration present) appears to be located downstream of glutamate receptors but upstream of PKA itself. It is possible to hypothesize that it is located in the post-synaptic density (PSD) assembly (Figure 8). The PSD is the network of scaffold proteins, receptors, enzymes and kinases that is located in close proximity to the post-synaptic membrane of excitatory synapses [62] and appears as an electron-dense structure in electron microscopy [63]. At least four major scaffold protein constituents of the PSD are affected in Ia synapses of mSOD1 MNs, namely Shank, Homer [22], PSD-95 [24] and AKAP (present work). The proper assembly of the PSD is necessary for the synaptic stabilization of glutamate receptors of several types [64, 65] and PSD disassembly is associated with the loss of synaptic glutamate receptor clusters [66, 67]. Furthermore, improper architecture of the PSD and in particular of AKAP reduces the efficiency of PKA activation in synapses [68], the phosphorylation of receptors by PKA and their stabilization [69], and synapto-nuclear signaling [70]. Finally, disturbances of PSD proteins and altered PKA signaling have been observed in non-SOD1 models such as cultured iPSC-derived MN with C9ORF72 mutations [25, 61] as well as in proteomic studies of human tissue [71].

**Figure 8:**
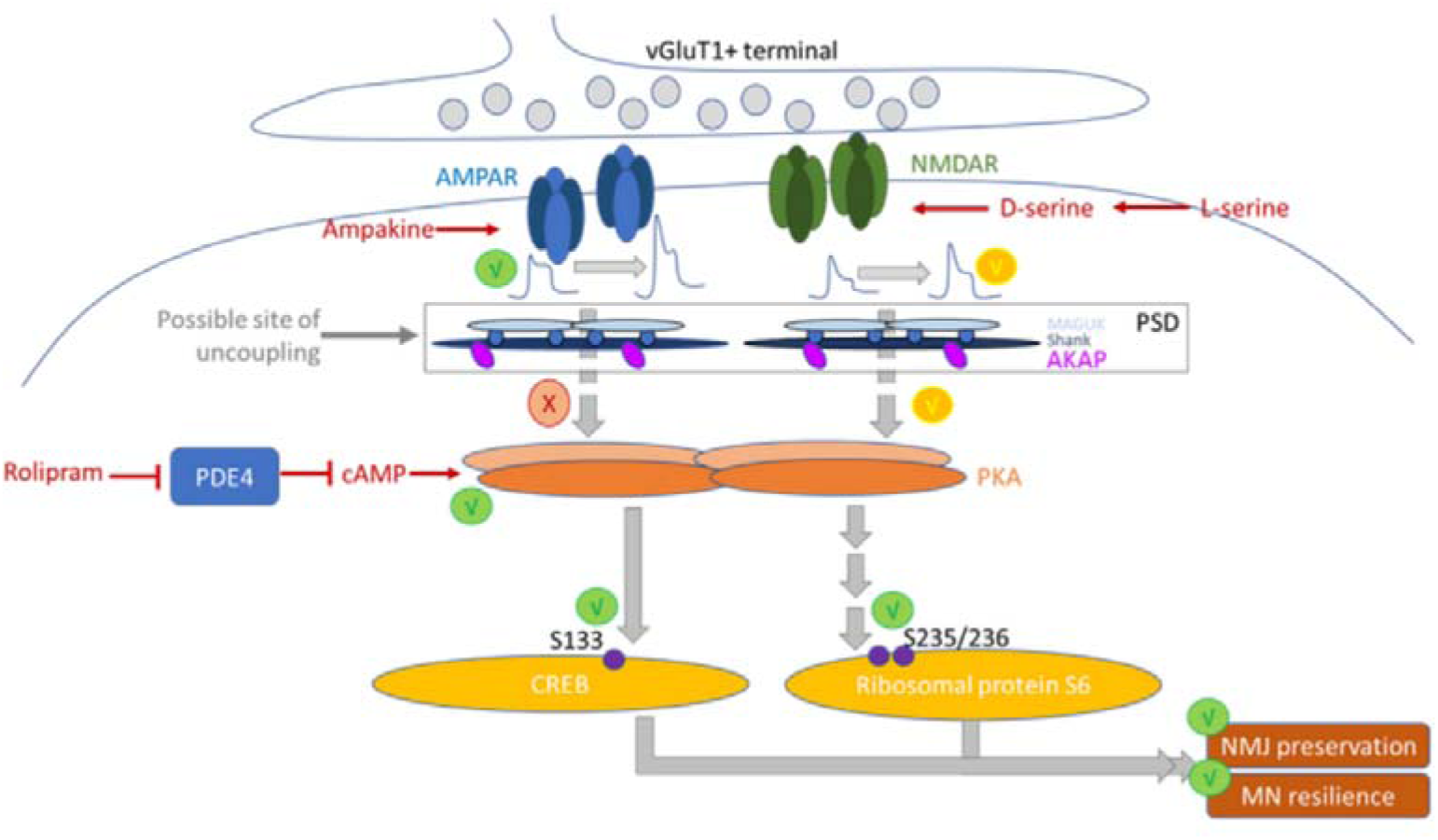
Possible site for synapto-nuclear signaling. Ampakine treatment and L-Serine/D-Serine treatment (trend only) are successful in increasing the EPSP size but fail at re-engaging the downstream signaling (L-Serine has a partial effect). On the other hand, directly increasing cAMP levels is sufficient to re-instate PKA signaling, as detected by the phosphorylation levels of the targets CREB and S6, indicating that the signaling cascade per se is functional if properly activated and can deliver neuroprotection. The site of uncoupling may be the post-synaptic density (PSD), the scaffold protein arrangement where signaling is activated. Disturbances in PSD members have been detected in ALS MN, including the loss of the PKA-scaffold AKAP.

### Restoring synapto-nuclear uncoupling has beneficial effects on ALS cellular hallmarks

We showed here that Rolipram, an inhibitor of the phosphodiesterase-4 which raises cAMP levels by preventing its degradation, can restore synapto-nuclear coupling. This is likely because the larger cAMP concentration boosts the PKA activity and thereby CREB phosphorylation. This is true not only during vibration-induced Ia activation, but also in a naturalistic environment (in which Ia are activated by the locomotor activity of the mice): in fact, ten-day delivery of Rolipram reduces the amount of misfolded SOD1 protein in the cytoplasm of MNs and the proportion of partially denervated neuromuscular junctions in presymptomatic mSOD1 mice. These results show that boosting the cAMP/PKA pathway restores synapto-nuclear coupling and, possibly in cooperation with other pathways modulated by cAMP, contributes to improving ALS hallmarks. These results are in line with the previously reported beneficial effect of ten-day activation of a DREADD (G)s on misfSOD1, LC3A, p62 [22]. Taken together these findings point toward a neuronal action of Rolipram (in contrast to an effect of Rolipram on non-neuronal cells, given the ubiquitous expression of PKA cascade components). Our results are in keeping with data in cortical-like neurons derived-IPSC from c9orf72 patients showing that a Rolipram treatment restores the PKA imbalance of regulatory and catalytic subunits, improves the dendritic arborization and the post-synaptic proteins (Jean-Gregoire et al. 2024). Importantly, the Rolipram treatment bypasses the adaptation mechanism (through desensitization/downregulation) that affects GPCR coupled with Gs [72], identifying PDEs as a more promising pharmacological target.

### Limitations of the study

This work has two unavoidable limitations. First, our investigation is based on the SOD1(G93A) mouse model of ALS; mutations in the SOD1 gene account for approx. 20% of familial cases of ALS, and these cases do not display a typical histopathological picture (i.e., they have no TDP-43 inclusions). Although we point to previous evidence from iPSC-derived motoneurons and human pathology leading toward the relevance of the findings beyond the SOD1 mutation [25, 61], this would require direct ascertainment. However, no mouse model with a profile of progression rate, phenotypic severity and reliability is currently available. Second, pharmacological agents are not cell-restricted in their action, and therefore, even if the system under study involves only Ia afferent and MNs, the small-molecules used (Rolipram, NBQX, L-serine) may also affect other neurons and non-neuronal cells (e.g., microglia) which may contribute to the final effect. Future work with genetically-targeted populations may elucidate the relative contribution of each cell type.

## Conclusion

Altogether, we demonstrated a functional uncoupling of activity-dependent synapto-nuclear signaling in MNs of mSOD1 mice. Enhancing cAMP/PKA signaling with phosphodiesterase-4 inhibitors partially restores synapto-nuclear coupling and mitigates crucial ALS hallmarks in MNs of presymptomatic ALS mice. These results implicate activity-dependent synapto-nuclear signaling as a contributor to MN resilience.

## Materials and methods

### Animals

This study used a total of 90 male B6SJL-Tg(SOD1*G93A)1Gur/J mice (hereafter, mSOD1) and 90 male B6SJL WT mice (hereafter, WT) bred at the Wielkopolska Center of Advanced Technologies at the Adam Mickiewicz University (Poznań, Poland) and the Animal House of Université Paris Cité (Biomed Tech *Facilities*). In Poznań, mice were housed in two per cage at the Poznań University of Physical Education Animal Facility (Poznań, Poland) with unlimited access to food and water. The animal room was set to a reverse light/dark cycle (12 h/12 h) with humidity and temperature maintained at 55 ± 10% and 22 ± 2 °C, respectively. In Paris, mice were grouped ≤5 with access to food and water *ad libitum* and housed in disposable and ventilated cages under a 12h/12h light/dark cycle with 40÷60% relative humidity.

mSOD1 mice display a phenotype similar to human ALS symptoms, with progressive limb muscle paralysis starting around postnatal day 90 (P90) and reaching the end-stage ALS around P120 [73]. Experiments were performed on mSOD1 mice between postnatal P45 - P55 which is considered a presymptomatic stage just before fast-fatigable MN denervation [74, 75]. Following European Union recommendations in Poznań, Ulm and Paris, our humane endpoints were defined as an inability of the mouse to reach food or water, the loss of more than 30% of body weight over 72 h, or an inability to rise or ambulate. None of the animals used in this study reached any of the humane endpoints. All procedures performed in this study were approved by the Poznań Local Ethical Committee (approval number 44/2018; Poznań, Poland); by the Regierungspraesidium Tubingen with licences no. 1404 and 1440; and by the Paris Descartes University ethics committee (CEEA34) and authorized by the french ministry for higher education and research (authorization number APAFIS#16338-2018052100307589). All authors held valid permits for working with laboratory animals and were appropriately trained in all experimental procedures.

### Retrograde labeling of motoneuron pool

Respective MNs were retrogradely labeled five days before the experiment with cholera toxin subunit b (CTb) conjugated with Alexa Fluor 555 (solution of 1.0 mg/ml CTb-555 with clean PBS) injected to MG and LG muscles (TS muscle group) of both right and left limb. The animals were anesthetized with 2-3% isoflurane in oxygen with isoflurane via a facemask. A bilateral incision was made on the skin of the hindlimb just above the TS muscle and 3 µl and 2 µl CTb-555 were injected into the left and right LG and MG muscles respectively with a Hamilton microsyringe with a 22-gauge needle attached. Following the injection, the needle was removed, the skin was sutured with sterile ligatures, and a subcutaneous injection of the analgesic and anti-inflammatory drug Meloxicam (0.4 mg/kg body weight) was given.

### Vibration of Achilles tendon

To induce activity-dependent metabolic pathways in TS MNs, an unilateral vibration of the Achilles tendon was performed. The mouse was initially injected with premedication (atropine 0.20 mg/kg s.c., Polfa and methylprednisolone 0.05 mg, s.c., Pfizer) and 15 minutes after anesthetized with an intraperitoneal injection of a drug cocktail (fentanyl 6.25 µg/ml (Polfa), midazolam 2.5 mg/ml (Polfa), medetomidine 0.125 mg/ml (Cp-Pharma) injected at 10 ml/kg body weight i.p. The depth of anesthesia was determined by the lack of hind limb withdrawal reflex. Subcutaneous needle ECG electrodes were positioned for heart rate monitoring, and the central temperature was maintained at 37 °C with an infrared heating lamp and an electric blanket (TCAT-2DF, Physiotemp). Then, an incision was made on the skin of the hindlimb just above the TS muscle in the right leg. The triceps nerve (composed of the nerves innervating the MG and LG muscles) was separated from the surrounding tissues and the common peroneal, sural, and distal tibial nerves were cut. The Achilles tendon was disinserted and firmly tied at the vibrator arm of a high-speed length controller (Aurora Scientific 322C, USA). Before the tendon was cut, the mouse paw was dorsiflexed to the maximum flexion angle (to indicate the muscle’s maximum physiological length, and following the cut, the tendon was preset at this length. The vibrator arm was sinusoidally driven with a frequency of 400Hz and peak-to-peak amplitude of 0.6mm. The duration of vibration was 0.2s with 1.8s interpulse interval. For the immunohistochemical analysis, the vibration cycle was repeated for 20 minutes. For electrophysiological investigations, each recorded MN received at least 10 vibration cycles.

### Preparation for electrophysiology

The methodology of electrophysiological investigation of Ia synaptic excitation on spinal MNs in the mouse has been previously described in detail in [22]. Initial steps of the surgery involved premedication, anesthesia as well as ECG and animal temperature maintenance, similar as described above for Achilles tendon vibration procedure. Next, a tracheotomy was performed, and the mouse was artificially ventilated with pure oxygen (SAR-1000 ventilator; CWE) with parameters adjusted to maintain the tidal CO_2_ level between 3 and 5% (MicroCapstar; CWE). Right and left external jugular veins were catheterized for administration of additional doses of the anesthetic cocktail (fentanyl 6.25 µg/ml (Polfa), midazolam 2.5 mg/ml (Polfa), medetomidine 0.125 mg/ml (Cp-Pharma); injected at 1.7 ml/kg body weight, i.v.), which were supplemented every 20-30 min; and for infusion of physiological buffer (4% glucose solution containing 1% NaHCO_3_ and 14% gelatine (Tetraspan; Braun) at 60 μl/h. Two pairs of horizontal bars (Cunningham Spinal Adaptor; Stoelting) were used to immobilize the vertebral column between the T13 and L2, and a laminectomy was made at the T13 - L1 vertebrae. The dura mater was removed from the exposed L3-L4 spinal segments to allow a glass microelectrode to be inserted into the spinal cord. Finally, the TS nerve was prepared for stimulation and the Achilles tendon was prepared for vibration, similarly as described above. The exposed tissues were covered with mineral oil in pools created by dissected and stretched skin flaps. At the end of the surgery, animals were paralyzed with pancuronium bromide (Pancuronium; Polfa; the initial bolus was 0.1 mg, followed by additional doses of 0.01 mg every 30-40 min). From this point on, each additional dose of the anesthetics was given at the same frequency as during the surgery or when ECG and/or PCO_2_ levels neared the maximal physiological values.

### Stimulation and recording

MN electrophysiological properties were measured intracellularly with glass micropipettes (tip diameter 1.0-1.5 μm, impedance 20-30 MΩ) filled with 2 M K-acetate for testing whether spikes were fired or not during the vibration-induced EPSPS (Supplemental Figure 1). In all other cases the microelectrodes were filled with a mix of 2 M K-acetate and 0.1 M QX-314 - a sodium channel blocker. The blocker was used to prevent the rare orthodromic action potentials in order to measure the amplitude of the EPSP at its peak. Intracellular recordings were obtained with an Axoclamp 900A amplifier (Molecular Devices) connected to a Power1401 interface (CED, sampling rate 20 kHz) operated by Spike2 software (CED). The amplifier system was used in the bridge mode to record vibration induced excitatory postsynaptic potentials (EPSPs) and cells resting membrane potential (RMP) or in discontinuous current clamp (DCC) mode (switching rate 7-8 kHz) to record the MN responses to square pulses of current necessary to determine the input resistance. Peripheral stimulation of the TS nerve with constant current pulses of 0.1 ms duration and amplitude up to 50 µA delivered at 3 Hz (DS4, Digitimer), but not more than 2 x threshold (Th) for the most excitable fibers in the nerve enabled us to identify motor neurons based on an “all-or-nothing” antidromic action potential. Importantly, this method of identification was still possible under the sodium channel blockade, as it takes QX-314 several seconds to diffuse from the recording electrodes and block the action potentials. An electrode positioned on the dorsal surface of the spinal cord was used to record the group I afferent volley in response to TS vibration. All the MNs selected for the analysis were characterized by a resting membrane potential (RMP) more hyperpolarized than −50 mV and an initial overshooting action potential. The TS spindle primary endings (from which Ia afferents originate) were activated by standardized Achilles tendon vibration protocol (described above) in order to elicit compound Ia EPSPs in homonymous MN. For each MN at least 10 vibration cycles were performed, and the resulting voltage traces were averaged. The maximal EPSP amplitudes were measured from the RMP (just before the onset of the vibration cycle) to the most depolarized part of the voltage response. The MN input resistance at RMP (RIN) was calculated by measuring the membrane voltage deflection in response to a series of small-amplitude square current pulses (−2 to +2 nA, 500 ms), as described in detail in [76]. At the end of the experiment, animals were euthanized with a lethal intravenous dose of sodium pentobarbital (200 mg/kg).

### Animal tissue processing and immunohistology

Tissue samples (spinal cord, muscles) were obtained at the Poznan site and, for independent experiments, at the Ulm site. At the Poznan site, mice were terminally anesthetized with a lethal dose of Morbital (200 mg pentobarbital/kg body weight, i.p.); at the Ulm site, mice were terminally anesthetized with Ketamine/Xylazine (100 mg/kg body weight ketamine and 16 mg/kg body weight xylazine, i.p.). At both sites, mice were then perfused transcardially (flow 6.5 ml/min) with 2ml/g 0.1 M phosphate-buffered saline (PBS) followed by 2.5ml/g ice-cold 4% paraformaldehyde (PFA). Spinal cord was dissected and post-fixed in PFA for 18h at 4°C and then cryoprotected in 30% sucrose in 0.1 M PB. Muscle samples (Tibialis Anterior and Gastrocnemius) were dissected and post-fixed in PFA 4% for 2h at 4°C before cryoprotection. The lumbar segment of the spinal cord was embedded in Tissue-Tek® OCT and frozen on dry ice. Transverse 40 μm thick sections were cut in the cryostat (Thermo Scientific Microm HM 520) at −19°C, washed in PBS, and used for immunostaining.

The immunofluorescence labelling of spinal cord sections was performed according to established methods previously reported [77]; the procedure was identical in Ulm and Poznan. Briefly, 40µm spinal cord cryosections (8-12 sections per animal, from L3-L5 lumbar segments, identified according to macroscopic morphology and anatomical landmarks) were washed in PBS-T (PBS+0.2% Triton X-100) for 1h and then transferred to blocking buffer (PBS + 0.2% Triton X-100, 3% BSA) for 2h at 24°C temperature on orbital shaker. Then the sections were manually transferred to new wells containing the primary antibodies mix (always including VAChT for MN identification plus additional antibodies as appropriate; see table 1) diluted in blocking buffer; the incubation was performed at 4°C for 40h on orbital shaker, followed by 4 x 30min washing steps in PBS-T. Secondary antibodies of appropriate specificity were diluted 1:500 in blocking buffer (see table 2); sections were incubated with secondary antibodies for 2h at 24°C, followed by 4 x 30min washing steps in PBS-T. Sections were manually transferred to glass slides for microscopy and mounted using Prolong-Antifade Gold mounting medium; the mounting medium was air-dried at 24°C for 18-20h before imaging. Muscle samples were sectioned along the longitudinal (insertion-tendon) axis at 15 µm thickness; sections were deposited directly on a glass slide for microscopy (Super-Frost™) and all following procedures were performed on adherent sections. Glass slides carrying the muscle sections were immersed in PBS-T 3 x 10min, and blocking/permeabilization was performed using PBS+ 2.5% Triton X-100 and 3% Donkey Serum. Sections were then covered with the blocking medium in which Alexa-647-conjugated Bungarotoxin (1:1000) and anti-synaptophysin (Synaptic Systems, 1:150) were added and incubated at 4°C for 40h; muscle sections were thereafter washed 3 x 20min in PBS-T, incubated with secondary antibody (Alexa488-anti mouse secondary, 1:500) for 2h, washed again and mounted in Prolong-Antifade Gold.

**Table 1.**
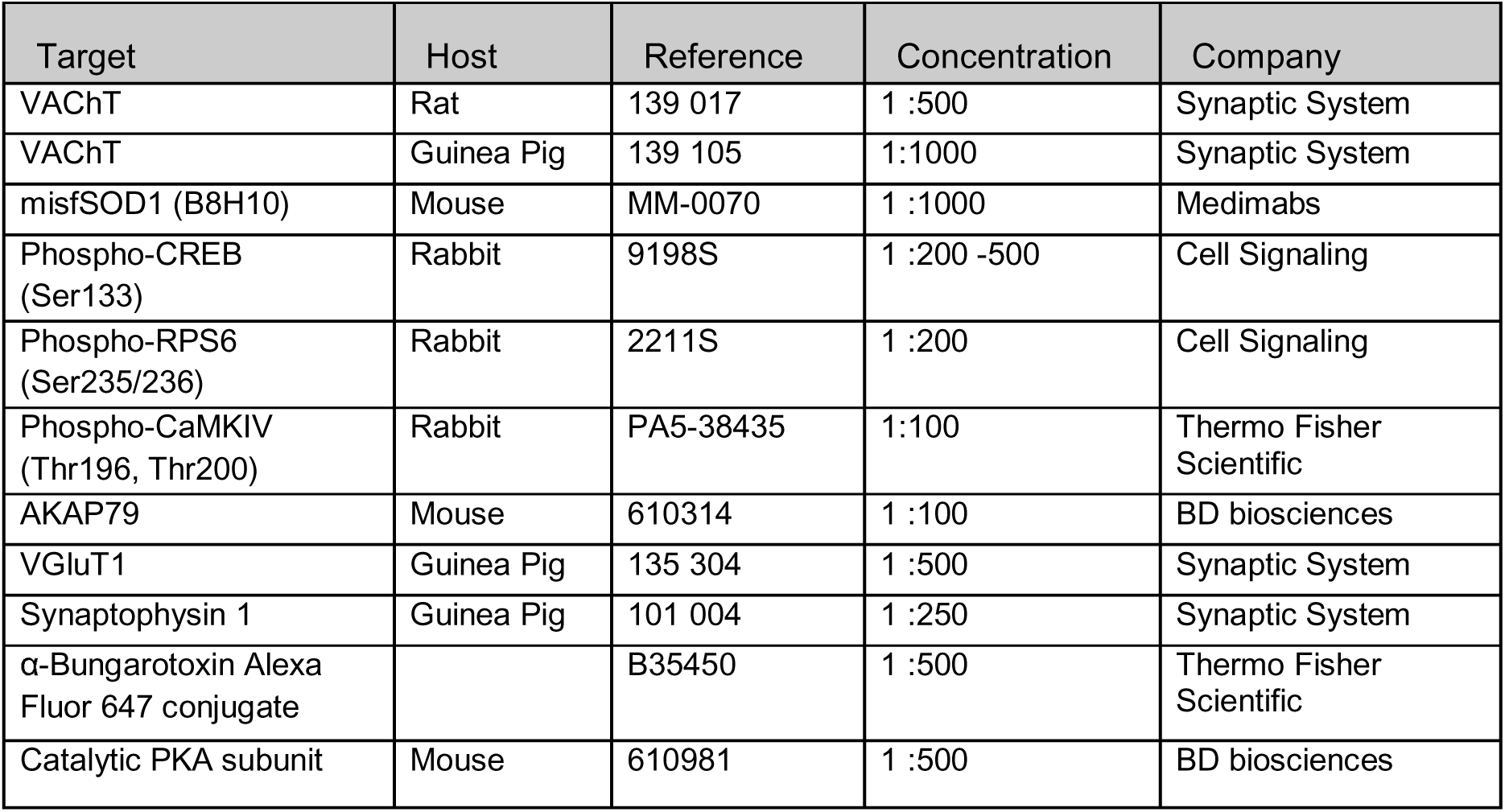
Primary antibodies.

**Table 2.** Secondary antibodies.

| Target | Host | Reference | Concentration | Company |
| --- | --- | --- | --- | --- |
| Anti-Rat Alexa Fluor 405 | Donkey | A48268 | 1:500 | Thermo Fisher Scientific |
| Anti-Goat Alexa Fluor 488 | Donkey | A11055 | 1:500 | Thermo Fisher Scientific |
| Anti-Goat CF 647 | Donkey | 20048 | 1:500 | Biotium |
| Anti-Mouse Alexa Fluor 647 | Donkey | A31571 | 1:500 | Thermo Fisher Scientific |
| Anti-Mouse Alexa Fluor 568 | Donkey | A10037 | 1:500 | Thermo Fisher Scientific |
| Anti-Rabbit Alexa Fluor 568 | Donkey | A10042 | 1:500 | Thermo Fisher Scientific |
| Anti-Rabbit Alexa Fluor 488 | Donkey | A21206 | 1:500 | Thermo Fisher Scientific |
| Anti-Guinea Pig Alexa Fluor 488 | Goat | A11073 | 1:500 | Thermo Fisher Scientific |
| Anti-Guinea Pig CF 405S | Donkey | 20356 | 1:500 | Biotium |

### Imaging in mouse spinal sections

Imaging was performed using a Leica DMi8 confocal microscope, equipped with 20X oil objective (for misfolded SOD, pCREB, pS6, CaMKIV, PKA) or 63X oil objective (for NR1, mGluR5, AKAP and synaptic PKA imaging). For each section, an artifact-free field of view corresponding to the ventral spinal cord was identified. Imaging was performed in non-saturating conditions using a 1µm optical sectioning thickness and laser power <5% of maximal output. For each field of view, a preliminary test of homogenous penetration of the antibody immunostaining was performed, verifying that the intensity profile was comparable at every depth. In sections devoid of technical artifacts and with homogeneous intensity profile, imaging was performed black-to-black, resulting in 25-30 individual optical sections (note that because of some dehydration during mounting, section thickness shrinks down from the nominal 40µm). Image files were recorded at 12-bits, with an intensity range between 0 and 4095; microscope settings were adjusted so that the majority of image elements had intensity between 500 and 3500, in the linear range of the detectors. Within each imaging session, samples from all treatments were imaged, to guarantee an homogeneous representation within imaging batches.

Muscle samples were imaged using an epifluorescence Keyence microscope equipped with a 20x air objective; exposure time was set on WT samples and kept constant across acquisitions. Alpha-bungarotoxin and synaptophysin 1 exposure times were set at 1/70 seconds and 1/10 seconds respectively. Multiple fields were acquired and stitched for overview acquisition.

### Image quantification in mouse spinal sections

Image analysis was performed in ImageJ. For the quantification of fluorescence intensity, the top and bottom 2-3 optical sections were excluded to avoid including cutting artifacts, and the remaining stack was flattened using a maximum-intensity projection algorithm. The resulting collapsed stack was subject to background subtraction using the rolling-ball option (50µm diameter) and then smoothed using a gaussian filter. On the resulting multi-channel image, MN were identified as cells in the ventral horn endowed with positive cytoplasmic VAChT surrounded by intensely VAChT-positive C-boutons (gamma-MN do not display C-boutons) (and positive cytoplasmic CTb signal in experiments with CTb tracer injection). Cell segments for which the negative image of the nucleus could not be detected were not considered as “cells” and not further analyzed; thus, bona-fide MN were considered when the imaging plane crossed the main diameter of the cell body. Before quantification, a quality control check was performed, and sections with holes or tears, antibody precipitation artifacts or foreign objects (such as grains of dust) were excluded. MN contour was manually traced in the VAChT channel, and the intensity of the other channels was then obtained sequentially using the fluorescence intensity option in ImageJ. For each image, the intensity of 4-6 regions of interest located away from the MN was also recorded, to further verify that the intensity range of all images was comparable and no batch-correction was necessary. For each animal, 10-12 artifacts-free ventral horn ROI were acquired and every MN in each ventral horn was quantified.

The quantification of neuromuscular junctions (NMJ) innervation was performed as previously reported [31], using a 70% colocalization threshold to define completely-innervated from non-completely innervated NMJ. At least 80 NMJ per animal were considered in correspondence of the segments L1-L2 of the TS [30] and at least 40 NMJs were considered from the TA.

### Human tissue processing, immunohistology and quantification

Human postmortem motor cortex paraffin sections were obtained from TargetALS biorepository (Table 3) and treated as previously described [61]. Briefly, sections were de-paraffinized in xylene and rehydrated in grading concentrations of ethanol. Sections were treated in 10mM citric acid pH 6 and 0.01% Triton X-100 at 100°C for 20 minutes before blocking in 1% BSA, 5% FBS, and 0.01% Triton X-100 in 1x PBS. Primary antibodies (PKA, BD Biosciences 610981; pCREB, Cell Signaling 9198S; MAP2, Thermo Fisher Scientific PA1-16751) were incubated in blocking buffer overnight at 4°C, while secondary antibodies were incubated at room temperature for 2 hours. Slides were mounted onto a glass slide using Prolong Gold mounting medium (Thermo Fisher Scientific) and imaged using a widefield microscope (Leica DMi8 Thunder) equipped with a cooled CMOS camera (DFC9000 GTC). Images were acquired as Z-stacks (0.21 μm step size) using a x63 lens. Images were deconvolved (AutoQuant, Media Cybernetics) and analyzed using Fiji/ImageJ software. First, 3D stacks were compressed using the max projection algorithm. DAPI stain was used to define the nuclear region, while MAP2 determined the contour of the cell soma (not shown in Supplemental Figure 3 for the sake of simplicity). Mean fluorescence intensity was measured in both regions independently.

**Table 3.** List of postmortem brain tissues used in the study.

| Subject ID | Gender | Condition | SOD1 Mutations | Site of Onset | Age at Onset (yrs) | Age at Death (yrs) | PMI (hrs) |
| --- | --- | --- | --- | --- | --- | --- | --- |
| NEUGD887RT1 | Female | ALS | A5V | Upper Limb | 29 | 32 | 5.16 |
| NEUPN968LVX | Female | ALS | I114T | Upper Limb | 74 | 75 | 31.5 |
| NEUXF746MEU | Male | ALS | A5V | Upper Limb | 63 | 64 | 19.9 |
| NEUHV604HT4 | Male | ALS | I114T | Limb | 58 | 59 | 15.6 |
| NEUUT630LZN | N/A | ALS | I114T | N/A | N/A | N/A | N/A |
| NEUDH441CDB | Male | NNC | -- | -- | -- | 47 | 32 |
| NEUEM656LYE | Male | NNC | -- | -- | -- | 62 | 46 |
| NEUGV102NYT | Female | NNC | -- | -- | -- | 63 | 25 |
| NEUKH683KEU | Male | NNC | -- | -- | -- | 89 | 7 |
| NEUPK860JBR | Female | NNC | -- | -- | -- | 91 | 24 |
| NEUWD570BTK | Male | NNC | -- | -- | -- | 68 | 10 |
| NEUWH642VP4 | Male | NNC | -- | -- | -- | 89 | 7 |

NNC: non-neurological control; PMI: postmortem interval; N/A: not available

### Drug preparation and administration

Cholera toxin subunit b (CTb) was conjugated with Alexa Fluor 555 (solution of 1.0 mg/ml CTb- 555 with clean PBS) and injected i.m. to left and right MG and LG muscle (TS muscle group), 5 days before applying Achilles tendon vibration (for both electrophysiological and immunohistochemical analysis) CX516-Ampakine (Sigma-Aldrich # SML1191, USA) an AMPA receptor agonist, was diluted in sterile saline and injected i.p. at the dose of 40 mg/kg (10 μl/g of body weight), 50 min prior to starting the vibration protocol for immunohistochemical analysis, or injected i.v. at the dose of 40 mg/kg (10 μl/g of body weight), immediately after recording a subset of control MNs for electrophysiological experiments.

L-serine (Sigma-Aldrich # S4311, USA), a NMDAR receptor co-agonist, was diluted in sterile saline, and injected i.p. at the dose of 2.5g/kg (10 μl/g of body weight), 20 minutes prior to starting the vibration protocol for immunohistochemical analysis, or injected i.v. at the dose of 2.5 g/kg, immediately after recording a subset of control MNs for electrophysiological experiments.

Rolipram (Sigma-Aldrich # R6520, USA), the PDE4 inhibitor was dissolved in 2% DMSO in sterile saline and injected i.p. at the dose of 1mg/kg [78] (10 μl/g of body weight). For the acute intervention, the i.p. injection was 40 min prior to starting the vibration protocol, or injection was performed i.v. immediately after recording a subset of control MNs for electrophysiological experiments. For chronic treatment, the i.p. injections were performed for 10 consecutive days starting at P34-44 with a final injection given one day before electrophysiological and immunohistochemical investigations. 2% DMSO (Sigma-Aldrich # RNBH9966, USA) in sterile saline injected at 10 μl/g of body weight served as a control intervention (Vehicle).

### Statistics

Plots were created in Prism v.6.0.7 (GraphPad Software), and statistical analysis was performed using R-Studio 2024.04.02 (Posit Software, PBC) with adequate libraries. Statistical analysis was carried out using mixed-effects models. Both histological and electrophysiological data was collected from multiple motoneurons per animal and comparisons included repeated measurements within animals as well as comparisons across independent groups. Thus, to adequately account for this non-independence of observations and variation between animals, models were specified with random intercepts for each animal. In vibration experiments, the fluorescence intensity of the vibrated side was normalized to the corresponding contralateral non-vibrated side from the same slice. The electrophysiological and cluster area data was analysed as raw values. The rest of the histological data was normalized to the mean intensity of the control group in each experiment to account for possible between-experiments variability. Random slopes were considered for within-animal predictors; but did not improve model fit. First linear models were fitted using the lmer function from the lme4 package in R (version 4.5.0). Model assumptions were evaluated with DHARMA’s [79] quantile-deviation, distribution, dispersion and outlier tests. Additionally, residual quantile–quantile plots and histograms were visually inspected. If model assumptions were violated, alternative models were considered. First, the outcome variable was log-transformed. If violations persisted, a generalized linear mixed model with a Gamma distribution and log link was fit using glmmTMB [80]. Final models satisfied all assumptions.

Except for the comparison of the effect of vibration of CTb- and CTb+ motoneurons on protein fluorescence intensity in wild-type mice (Figure 1), all analyses involved only comparisons of outcome measures between two groups. For the analysis of the impact of unilateral Achilles tendon vibration on pCREB, pS6 and pCamKIV levels in WT (Figure 1), the fixed effects were the vibration side (nvib, vib) and CTb labelling (CTb+, CTb-) and both factors as well as their interaction effect were included as fixed effects. For the analysis of the impact of unilateral Achilles tendon vibration on pCREB, pS6 and pCamKIV levels in mSOD1 mice (Figure 2), the fixed effects were the vibration side (nvib, vib). For the analysis of acute impact of NMDA and AMPA receptors agonists on TS MNs electrophysiological profile (Figure 3), the fixed effect was the treatment (Before, After), and for the histological analysis, the fixed effect was the vibration side (nvib, vib). For the analysis of NMDA and PKA clusters, in WT and mSOD1 animals (Figure 4), the fixed effect was the genotype (WT, mSOD1). For the analysis of the impact of acute rolipram treatment on MNs electrophysiological profile and pCREB, pS6 and pCamKIV levels in mSOD1 animals (Figure 5), for electrophysiological data, the fixed effect was the treatment (Before, After), and for the histological analysis, the fixed effect was the vibration side (nvib, vib). For the analysis of chronic rolipram treatment on MNs electrophysiological profile and pCREB, pS6 levels (Figure 6), misfoldedSOD1 levels and partially innervated NMJ levels (Figure 7), the fixed effect was the treatment (Vehicle, Rolipram) for both electrophysiological and histological data. In the supplementary material, for the analysis of NMDA and mGluR5 receptors cluster organization (Supplementary figure 2), between WT and mSOD1 animals, the fixed effect was the genotype (WT, mSOD1). For human data, in the analysis of PKA and pCREB levels on cortical layer V neurons (Supplementary Figure 3), the fixed effect was the genotype (WT, mSOD1). For all analysis the random intercept was the animal except for the Supplementary Figure 3, where the fixed effect was the patient. Kenward-Roger method was used to adjust degrees of freedom. Results are reported as the average of normalized values ± SE, except for the electrophysiological and cluster area where the average of raw data ± SE is presented. The specific test used for each analysis, including potential log-transformation of the outcome variable, is detailed in the corresponding figure legend.

## Data availability

The datasets generated and/or analysed during the current study are available from the corresponding authors on request.

## Acknowledgments

The authors acknowledge the animal research and breeding facility and the imaging facility of BioMedTech Facilities at Université Paris Cité (INSERM US36 / CNRS UAR2009) for their support and expertise. We also thank the Poznan Supercomputing and Networking Center affiliated with the Institute of Bioorganic Chemistry of the Polish Academy of Science for providing storage space for the raw electrophysiological data repository.

## Author contributions

Conceptualization: DZ, FR, MB, GC; Data curation: KG, PZ, SMD, BW, MB; Formal analysis: KG, PZ, SMD, CJ, RS; Funding acquisition: DZ, FR, MB; Investigation: KG, PZ, BW, BO, OA, MB, CJ, RS; Methodology: DZ, FR, MB; Project administration: MB, DZ, FR; Resources: MB, DZ, FR, CF; Software: SMD, MB, PZ; Supervision: DZ, MB, FR, CF; Validation: DZ, FR, MB; Visualization: KG, PZ, BW, CJ; Writing-original draft: DZ, FR, MB, SMD, KG; Writing-review&editing: DZ, FR, MB, CF, GC, SMD, CJ, KG, PZ, BW.

## Ethical approval and consent to participate

All procedures performed in this study were approved by the Poznań Local Ethical Committee (approval number 44/2018; Poznań, Poland); by the Regierungspraesidium Tubingen with licences no. 1404 and 1440; and by the Paris Descartes University ethics committee (CEEA34) and authorized by the french ministry for higher education and research (authorization number APAFIS#16338-2018052100307589). All authors held valid permits for working with laboratory animals and were appropriately trained in all experimental procedures. Human postmortem motor cortex was obtained from TargetALS brain Bank collection. (Human Postmortem Tissue Core - Target ALS https://share.google/Atb2pchOlOkXP0QG6).

---

**Supplemental Figure 1:**
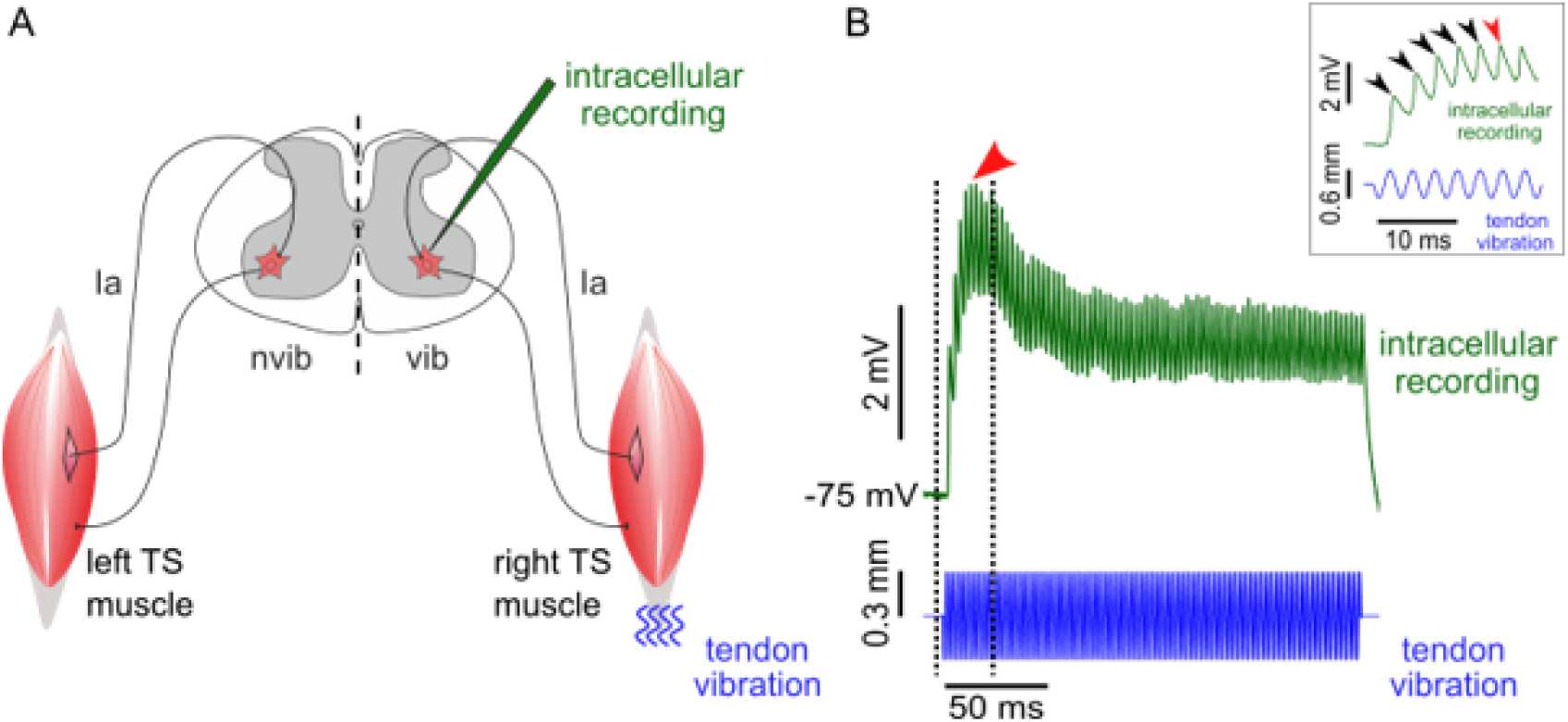
High-frequency vibrations of the Achilles tendon elicit EPSPs in TS motoneurons. **A,** experimental design for the intracellular recordings of EPSPs evoked by unilateral Achilles tendon vibration in TS MNs. **B,** representative example of a vibration-induced Ia EPSP in a TS motoneuron. Top trace (green): voltage response, bottom trace (blue): vibrator arm movement. The region between two dotted lines is magnified and shown in the insert on the top right of the panel. Each vibration cycle triggers a unitary Ia EPSP and temporal summation of successive EPSPs (insert, black arrows) creates the peak EPSP (red arrowheads) which then decreases and plateaus due to the deactivation of Ih current. TS - triceps surae muscle, Ia - Ia proprioceptive afferent, nvib - non-vibrated side, vib - vibrated side.\

**Supplemental Figure 2:**
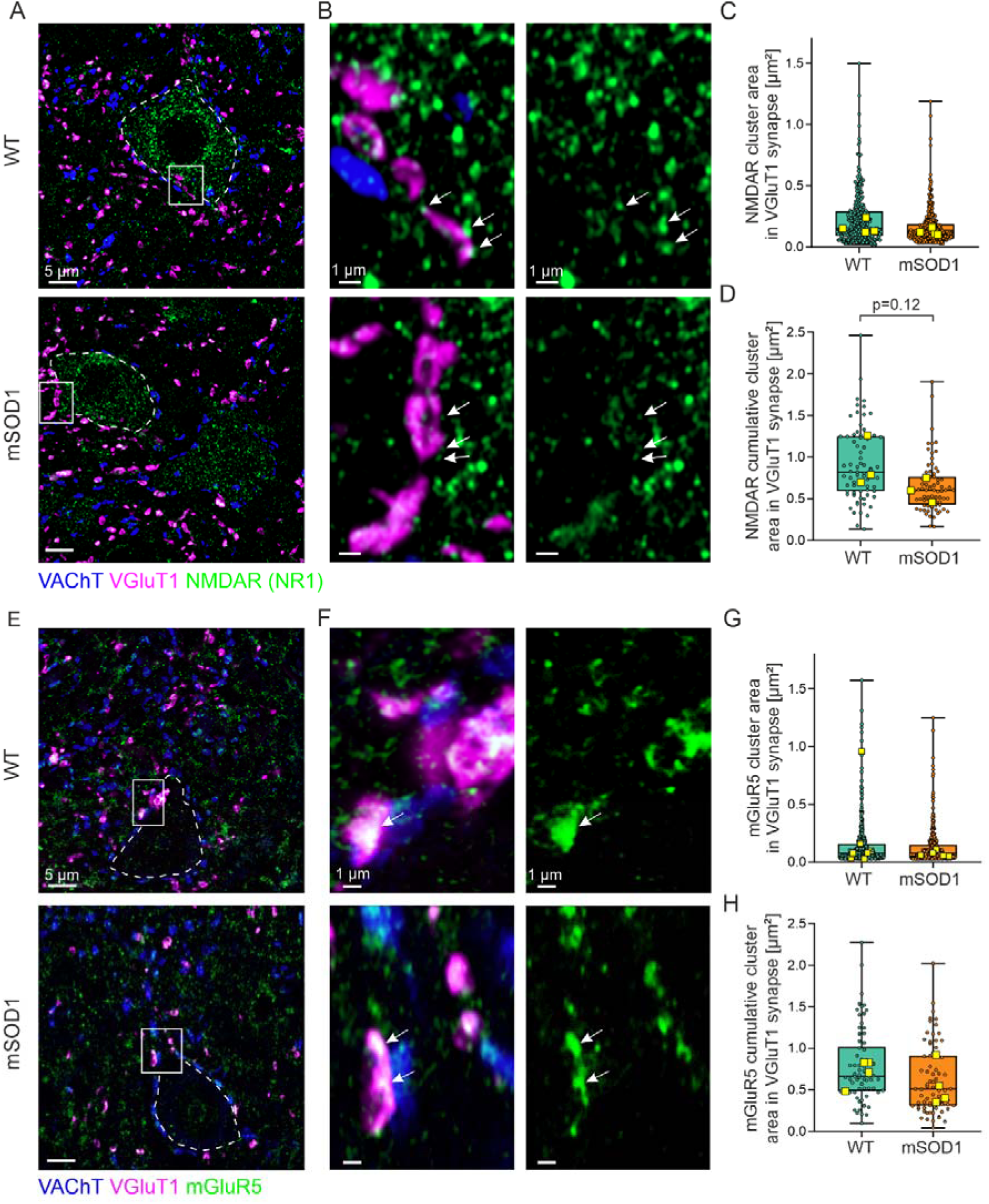
Representation of NMDA and mGluR5 receptors at VGluT1 synapses into MN from WT and mSOD1 mice. **A,** representative image of lumbar MN of WT (top) and mSOD1 (bottom) mice labelled for NMDA receptor clusters at the post-synaptic side of the VGluT1 synapse. **B,** magnification of the boxed region in A, showing a representative VGluT1 synapse, with white arrowheads pointing to the NMDAR clusters. **C,** boxplots displaying the distribution of normalized data for NMDAR cluster size and showing no difference between WT and mSOD1 mice (0.24±0.02 µm^2^ vs. 0.16±0.02 µm^2^, EMMs ratio=0.75±0.13, t_6_=-1.63, p=0.15, LMM on log-transform outcomes). **D,** similar as for C, but showing data distribution pointing to a tendency for decreased cumulative NMDAR cluster area in mSOD1 animals (0.97±0.10 µm^2^ vs. 0.69±0.08 µm^2^, EMMs ratio=0.72±0.13, t_5.99_=-1.81, p=0.12, LMM on log-transform outcomes, n=8 mice). **E-F,** similar as for **A-B**, but showing the mGluR5 subunits of the AMPA receptor at the post-synaptic side of the VGluT1 synapse as indicated by white arrowheads. **F - H**, similar as for **C-E**, but showing the lack of significant difference in mGluR5 clusters organization between WT and mSOD1 mice (0.16±0.02 µm^2^ vs. 0.13±0.03 µm^2^, EMMs ratio=1.00±0.13, t_5.98_=-0.017, p=0.99 for cluster size, LMM on log-transformed outcomes, and 0.90±0.14 µm^2^ vs. 0.64±0.11 µm^2^, EMMs ratio=0.71±0.17, t_5.95_=-1.440, p=0.20 for cumulative cluster size, on log-transformed outcomes, n=8 mice). For both **A** and **E**, the MN soma is indicated by white dashed lines as traced from VACht labelling. Scale bar 5 µm for **A** and **E**, and 1µm for **B** and **F**. Boxplots and statistical description as in Figure 1.

**Supplemental Figure 3:**
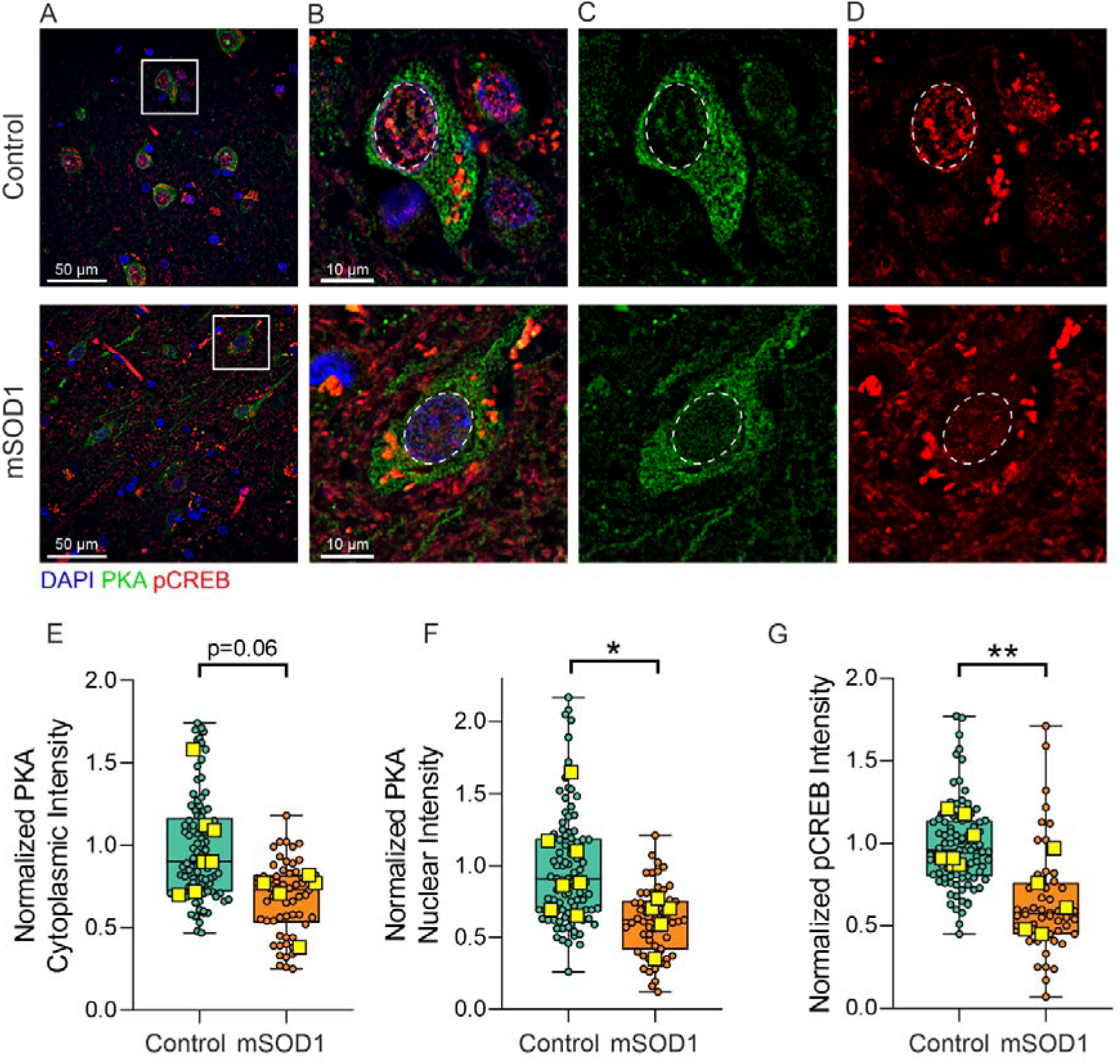
PKA signaling is disrupted in cortical motoneurons of ALS patients. **A**, Representative images of human cortical layer V neurons, harvested post-mortem from healthy donors (top row) and ALS patients harbouring the SOD1 mutation (bottom row). **B-D,** show magnification of the boxed areas from panel A, and display the baseline fluorescence intensity of PKA (**C**) and pCREB (**D**) signal, with superimposed labelling shown in **B**. **E-G,** boxplots displaying the normalized data distribution for lower cytoplasmic PKA (1.00±0.11 vs. 0.69±0.08, EMMs estimate=7067±3400, t_10_=2.053, p=0.06), nuclear PKA (1.00±0.13 vs. 0.62±0.07, EMMs estimate=6732±2990, t_10_=2.253, p=0.04, LMM) and pCREB (1.00±0.06 vs. 0.65±0.10, EMMs estimate=3194±953, t_10.2_=3.351, p<0.01, LMM) levels in SOD1 patients (n=4) when compared to healthy controls (n=7). Scale bar 50 µm for **A**, and 10 µm for **B-D**. In **B-D**, the MNs nuclei are indicated by white ovals guided by DAPI labeling. Box plots and statistical description as in Figure 1.

